# Patient-specific simulation of Retinal Hemangioblastoma provides new perspectives on the role of antiangiogenic therapy

**DOI:** 10.1101/2023.02.24.529937

**Authors:** Franco Pradelli, Giovanni Minervini, Silvio C.E. Tosatto

## Abstract

Retinal Hemangioblastoma (RH) is the most frequent manifestation of the von Hippel-Lindau syndrome (VHL), a rare disease associated with the germline mutation of the von Hippel-Lindau protein (pVHL). An emblematic feature of RH is the high vascularity, which is easily explained by the overexpression of angiogenic factors (AFs) arising from the pVHL impairment. The introduction of Optical Coherence Tomography Angiography (OCTA) allowed observing this feature with exceptional detail. However, our understanding of RH is limited by the absence of an animal model fully recapitulating the tumor. Here, we exploit a cancer mathematical model as an alternative way to explore RH development and angiogenesis. We derived our model from the agreed pathology for this tumor and compared our results with patient-specific OCTA images. Our simulations closely resemble the medical images, proving the capability of our model to recapitulate RH pathology. Our results also suggest that angiogenesis in RH occurs suddenly when the tumor reaches a critical mass, with full capillary invasion in the order of days. These findings open a new perspective on the critical role of time in antiangiogenic therapy in RH, which has resulted ineffective. Indeed, it might be that when RH is diagnosed, angiogenesis is already too advanced to be effectively targeted with this mean.

## INTRODUCTION

The von Hippel-Lindau syndrome (VHL) is a genetic disease predisposing to cancer and cysts development in multiple organs (Crespigio et al., 2018). Despite its limited incidence (1/36000 births), the study of this disease has led to crucial insights in cancer and biological research, the first being the characterization of the von Hippel-Lindau protein (pVHL) (Richard et al., 2013). Indeed, this protein, which is mutated in VHL, is now recognized as a central element of the cellular oxygen-sensing pathway and a key oncosuppressor (Gossage et al., 2014; Minervini et al., 2015). Among VHL-related tumors, Retinal Hemangioblastoma (RH) is the most frequent and earliest to occur (Karimi et al., 2020; Magee et al., 2009; Ruppert et al., 2019). It usually presents as a slowly-growing, vascular benign tumor but can lead to severe vision impairment (Magee et al., 2009). It is often associated with exudation and, at later stages, with retinal vessels enlargement and tortuosity (Figure S1). RH pathology has been debated in the past, but today the scientific community agrees on the essential steps of RH development. RH originates early in the patient’s life (Karimi et al., 2020), probably from progenitor cells arrested during development (Park & Chan, 2012). These cells lose heterozygosity in pVHL and differentiate into foamy tumor cells observed in RH and central nervous system hemangioblastoma (Chan et al., 2007). The mutation is linked to angiogenic factors (AFs) overexpression, such as the vascular endothelial growth factor (VEGF) and the platelet-derived growth factor (PDGF). Indeed, high VEGF concentrations have been reported in RH cells (Chan et al., 2007) and VHL patients’ vitreous (Los et al., 1997). Finally, AFs induce the formation of novel blood vessels (angiogenesis), explaining the highly vascular nature of RHs.

Even though several studies support RH pathology, it still lacks validation *in vivo* due to the absence of a reliable animal model (Park & Chan, 2012). A significant obstacle is that VHL impairment is lethal for mice embryos, but even conditional knockout could not recapitulate RH development (Wang et al., 2018). The absence of an animal model also limits our understanding of the effect of antiangiogenic therapy (AAT) in RH. Given the role of angiogenesis in RH, there were great expectations about the effectiveness of this therapy (Harris, 2000). Several studies have assessed the application of VEGF and PDGF inhibitors (Wiley et al., 2019), the latest being presented in 2021 (Hwang et al., 2021). However, the main observed effect was exudation reduction, with a minimal or absent decrease in tumor volume.

Animal models might not be the only way to explore RH development in time. Cancer Mathematical Models (CMMs) are proving to be an exceptional tool for cancer research, both from the biological and clinical perspectives. From one side, CMMs allow simulation of the interplay between complex biological phenomena (e.g., nutrients distribution, biochemical reactions, immune system), unveiling non-trivial aspects of tumor growth (Chauviere et al., 2010). On the other side, they can integrate and process patients-specific data (e.g., omics, imaging) to predict cancer evolution or treatment outcome, an approach known as “predictive medicine” or “precision medicine.”

There are different kinds of CMMs, exploiting different data and mathematical tools. Phase Field Models (PFMs) are based on partial differential equations (PDEs) describing the evolution in time of the boundaries of biological tissues, including neoplasms (Travasso, Castro, et al., 2011). These models can easily integrate with imaging data (Xu et al., 2020) and simulate cancer development at a tissue scale and in a patient-specific way (Lorenzo et al., 2016). Moreover, PFMs have yielded promising results in reproducing cancer development and cancer-related phenomena. Lorenzo G. and collaborators presented a prostate cancer simulation at an organ scale (Lorenzo et al., 2016) which allowed them to assess the effect of pressure on its development (Lorenzo et al., 2019). Travasso et al. developed a PFM to simulate angiogenesis (Travasso, Poiré, et al., 2011), which was later employed to study the effect of interstitial flow in tumor-induced angiogenesis (Moure et al., 2022). Xu J. et al. have integrated the same model with medical imaging to simulate tumor growth and vascularization integrating photoacoustic images (Xu et al., 2020).

RH represents an ideal case study for PFMs for several reasons, especially considering the coupling between tumor growth and angiogenesis. First, it is small, usually a few millimeters in size (Karimi et al., 2020). This reduces the computational effort to simulate the total tumor volume and the surrounding tissue. Second, Optical Coherence Tomography Angiography (OCTA) enables non-invasive access to patient-specific data for both the cancer borders and the capillaries (Figure S1). These images are an invaluable validation source for tumor-induced angiogenesis, especially compared to traditional RH imaging techniques (Sagar et al., 2018). Third, pVHL impairment decouples vascularization from hypoxia, slightly reducing the model complexity. Indeed, many CMMs have been employed to explore the formation of new blood vessels in cancer (Levine, Pamuk, Bölümü, et al., 2001; Phillips et al., 2020; Xu et al., 2016). However, since this event typically occurs in hypoxic conditions, these models must account for several variables. For instance, the oxygen distribution and the existence of a hypoxic core overexpressing AFs are often included in these CMMs, but they are both challenging to observe *in vivo*.

In this study, we use PFMs to simulate RH development and angiogenesis at the tumor scale and in a patient-specific way. We selected a case report showing an early-stage RH, including OCTA images of capillaries and tumor borders and used these data to develop a simple but comprehensive model in respect of the agreed pathology of RH. When possible, we derived the model’s parameters from experimental evidence. Finally, the selected case report (SCR) was used to validate our results and explore multiple scenarios through the variation of critical parameters.

## RESULTS

### Mathematical Model

Our PFM accounts for three main elements to describe RH dynamics, each represented by a scalar function: the tumoral mass 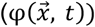, which equals 1 inside the tumor and 0 outside; the capillaries field 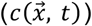, which has value ≈ 1 in the intravascular space and ≈ −1 outside; and the AFs 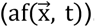. A schematic representation of the model is reported in Figure 1. In the notation we just used, t represents time and 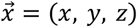 represents space. The plane defined by x and y is parallel to the retinal layers, and z perpendicular, pointing in the direction of the inner eye. Our model implies the definition of several parameters. See Table 1 for an overview of the parameters used and the Supplementary Information (SI) for an extensive discussion on their derivation.

**Table 1.**
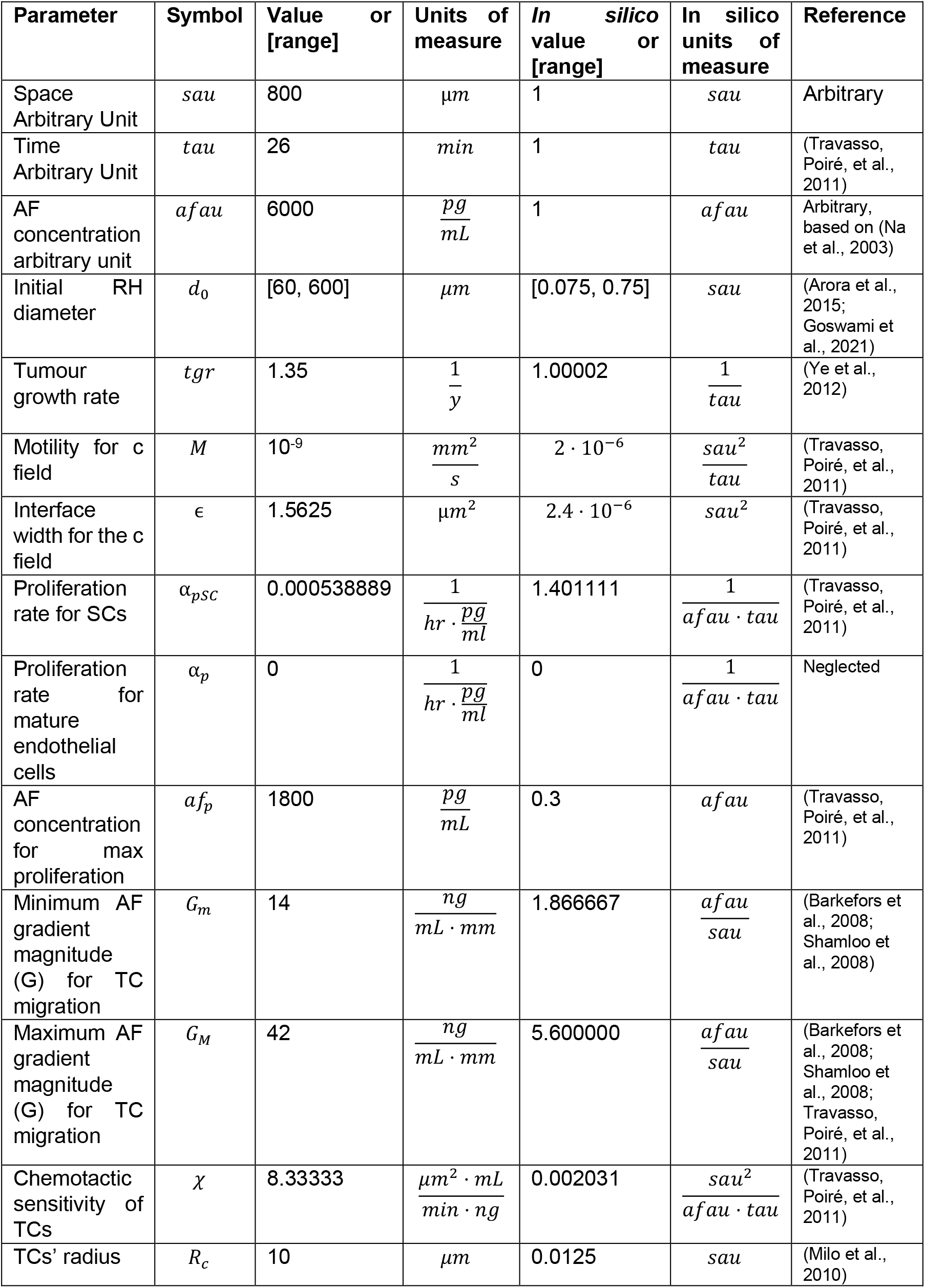
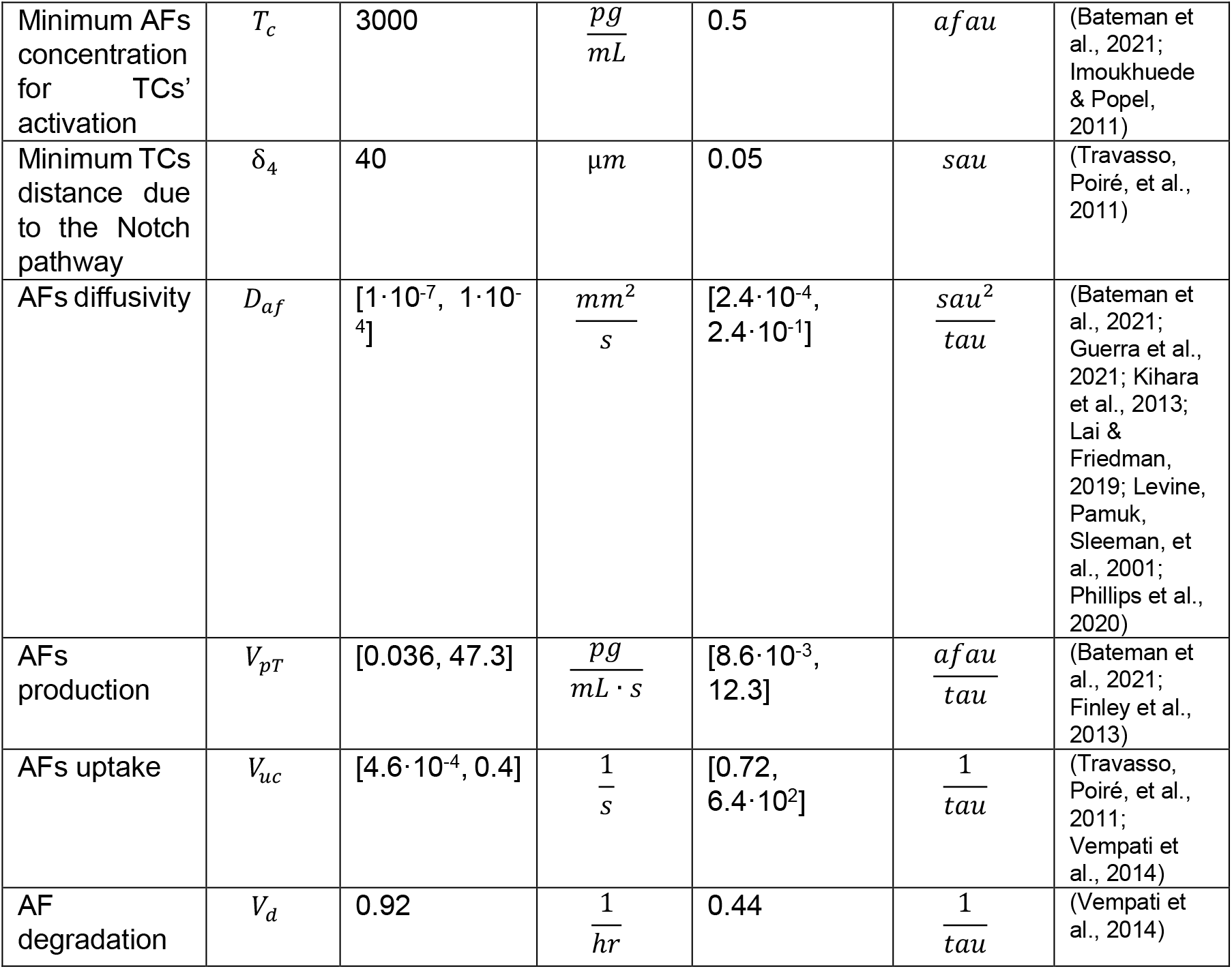
Parameters used in the simulations presented. When possible, we derived the values from experimental evidence, otherwise an estimation was used. For an extensive discussion on the derivation of each value, see Supplementary Information. When we employed multiple values for a single parameter, we provided a range.

**Figure 1.**
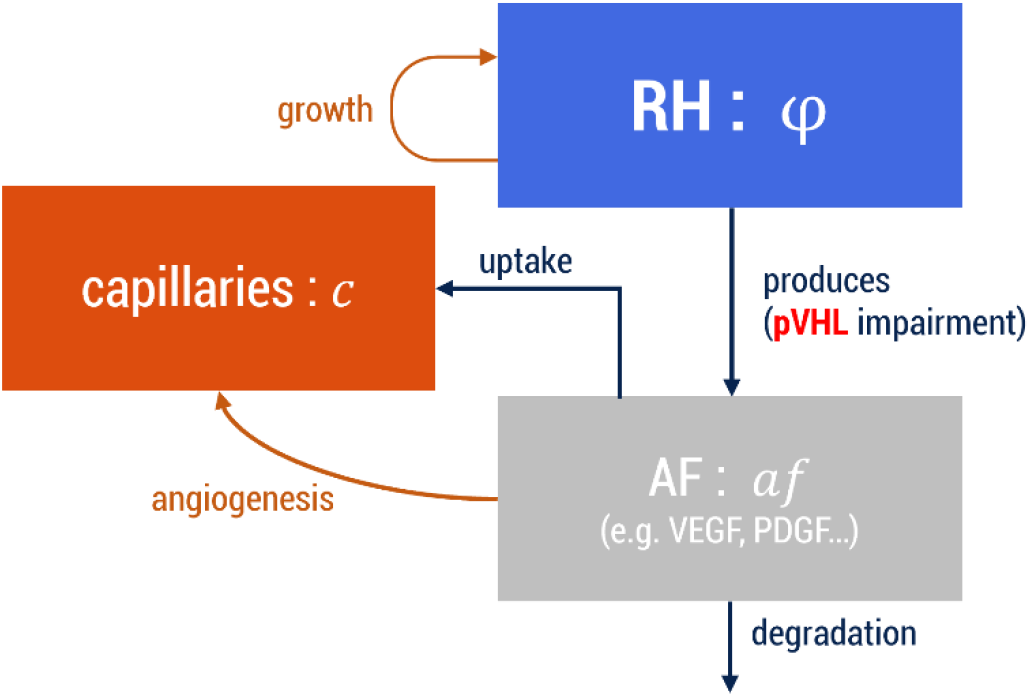
Schematic representation of the PFM we employed in our study. RH = Retinal Hemangioblastoma; AF = Angiogenic Factors. The arrows represent how the different elements interact with each other. For each entity, the name of the scalar field is also reported.

Since RH is often rounded-shaped, and the RH reported in SCR has an ellipsoidal shape (see Figure 2a), we defined φ as a slowly growing ellipsoid in the retina:

**Figure 2.**
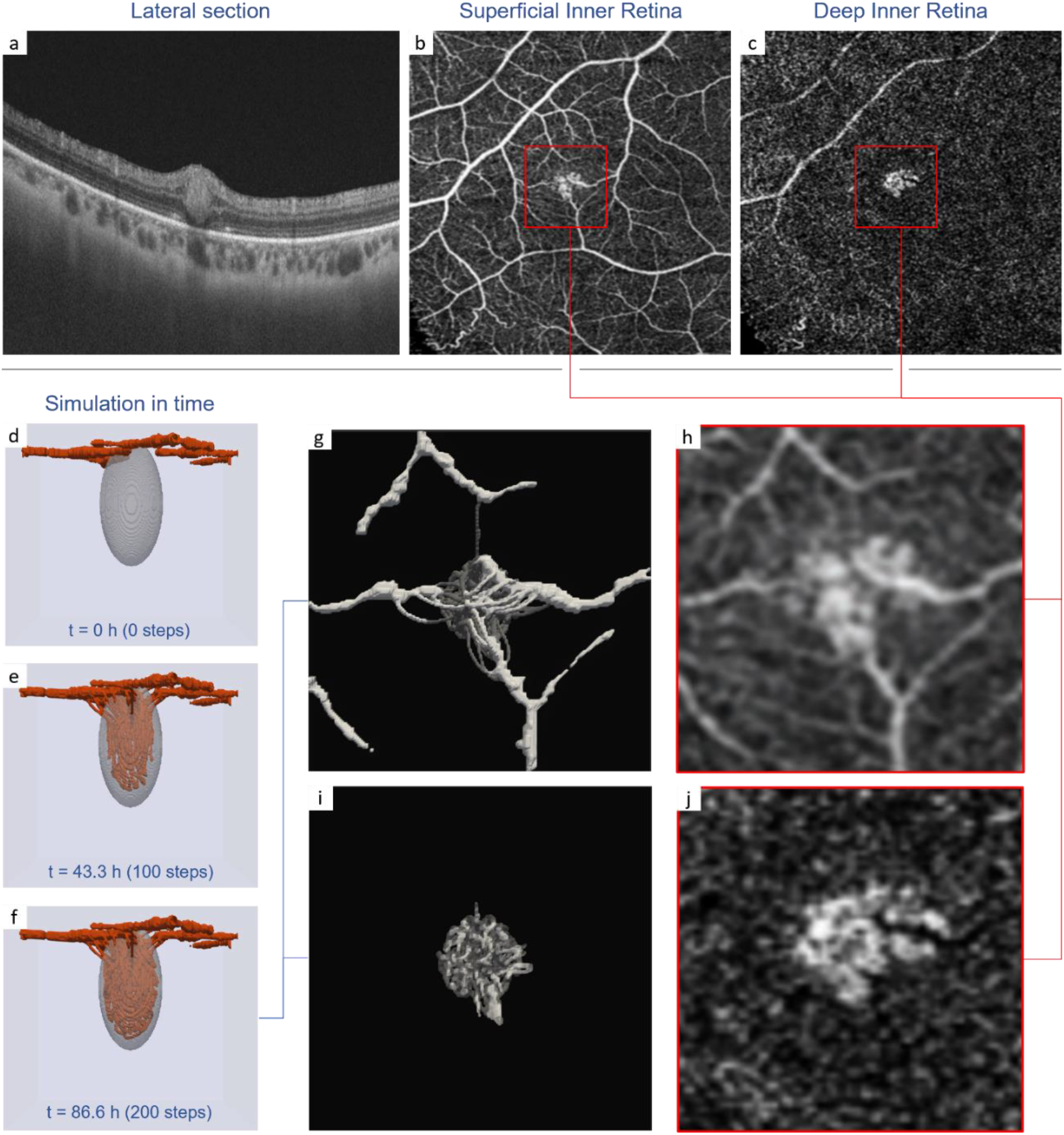
a)-c) OCTA images derived from the SCR (Goswami et al., 2021). The red square is the area corresponding to the simulation mesh. d)-f) evolution of the PFM in time. At t = 0 (d), it is possible to observe the initial avascular tumor, and the initial capillaries reconstructed in 3D. RH growth is almost unobservable, while tumor-induced angiogenesis leads to full tumor invasion in about 87 hours. g) top view of the simulation result compared to the actual capillaries observed in the patient (h). i) top view of the simulation result sectioned at the level of a deeper retinal layer, compared with the actual capillaries observed in the patient (j). The simulation was obtained setting *d*_0_ = 400 [*μm*], *D*_*af*_ = 4.24 · 10^−5^ [*mm*^2^ · *s*^−1^], *V*_*pT*_ = 47.3 [*pg* · *mL*^−1^ · *s*^−1^], *V*_*uc*_ = 4 · 10^−2^ [*s*^−1^].

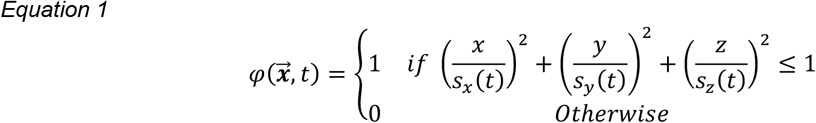

Where:

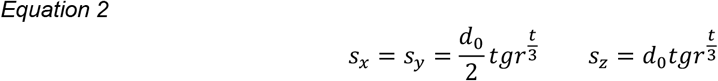

With *d*_0_ corresponding to the initial tumor diameter in the section parallel to the retinal layers and *tgr* corresponding to the tumor growth rate.

To model the capillaries’ dynamics, we used a hybrid model presented by Travasso and collaborators (Travasso, Poiré, et al., 2011), which merges a PFM and an agent-based approach. Coherently to the mainstream theory for angiogenesis, this model assumes that AFs induces novel capillaries activating tip cells (TCs), which direct the capillaries formation together with the stalk cells (SCs). We shortly explain this model in Materials and Methods and SI, but we refer to the original publication for an extensive presentation of the model. We introduced a minor variation to the model to better reproduce our case study, which is explained in Materials and Methods and SI. AFs dynamics is regulated by the following PDE:

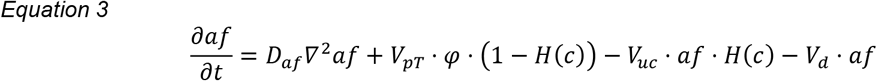

The first term (*D*_*a*f_∇^2^af) accounts for AFs’ diffusion. The second term (V_pT_ · φ · (1 − H(c))) accounts for AFs’ production, which, in line with the agreed pathology for RH, takes place inside the whole neoplasm. We further assumed that AFs production does not take place inside the capillaries. Thus, we added the term (1 − *H*(*c*)), where *H* is the Heaviside function. The third term (*V*_*uc*_ · *af* · *H*(*c*)) accounts for the uptake of the AFs from the capillaries. Since our model does not account for blood flow, this term encapsulates many complex phenomena: a) the uptake of AFs from the endothelial cells due to the binding with membrane receptors; b) the transport of AFs away due to blood flow circulation; c) the uptake of AFs from platelets, which can bind specific molecules such VEGF (Kut et al., 2007). Finally, the term (*V*_*d*_ · *af*) accounts for natural AF degradation.

To compute the initial condition, we assume that *af* distribution was near the equilibrium before sprouting angiogenesis. Thus, we solve <u>Equation 3</u> at equilibrium 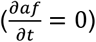 for a given initial tumor and capillaries’ network. As boundary conditions, we assume natural Neumann again:

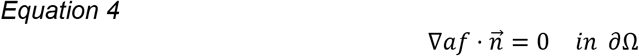

So, we assume there is no AF flux through the mesh boundaries. This might not be precisely the case, but it is still reasonable to assume that AFs flux through the boundaries is negligible compared with the local production by the tumor and uptake by the blood vessels.

## Simulation and comparison with OCTA

Using a putative initial capillary network derived from the SCR (see Materials and Methods) and several initial tumors, we employed our PFM to simulate RH development and angiogenesis. As shown in Figure 2, the result closely resembles the capillaries network observed in the patient’s OCTA images.

The simulation reported in Figure 2 is obtained setting *d*_0_ = 400 [μ*m*], *D*_*af*_ = 4.2 · 10^−5^ [*mm*^2^ · *s*^−1^], *V*_*pT*_ = 47.3 [pg · *mL*^−1^ · *s*^−1^], *V*_*uc*_ = 4 · 10^−2^ [*s*^−1^]. Since both *V*_*pT*_ and *V*_*uc*_ are in the high spectrum of the tested range (see Table 1), the simulated scenario is an RH characterized by high AF production and transport. The simulated tumor’s dimension is smaller than the tumor reported in SCR, which is about 600 [μ*m*]. Thus in the context of a growing mass, the simulated scenario represents a situation where tumor-induced vascularization was triggered before the tumor reached its maximum extension, which is quite realistic.

In 200 time steps (≈ 87 *h*), we observe that angiogenesis is quick enough to drive RH from the initial avascular state to complete tumor vascularization. At the same time, tumor growth at this stage is barely noticeable. Most of the capillaries’ network is already formed after the first 100 steps (≈ 43 h). Indeed, we observe that the number of TCs peaks at 100 steps and then steadily decreases in the last part of the simulation (see Figure S2).

Unsurprisingly, we observe that AFs concentration is higher inside and around RH, and that concentration decreases with vascularization (Figures S3, S4). Initial AFs concentration is initially in the range [1, 23] 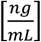 (average: 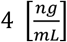), while at the end, the range shrinks to [0.5, 6] 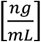 (average: 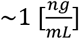).

Similarly, the AFs gradient (G) is stronger at the beginning and steadily decreases over time (Figures S3, S5). Initially, G ∈ [0.01, 152] 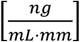 (average: 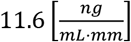), while at the end, the range reduces to [0.003, 40.7] 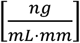 (average: 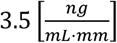). Notably, our simulation shows that the combination of production inside the tumor and uptake in the capillaries is sufficient to drive angiogenesis during the simulation.

### Effect of parameters on RH-induced angiogenesis

The simulation reported in the previous section recapitulates the observation of the SCR. However, parameters value always have a critical impact on CMMs. Thus, we explored the simulation results for different parameters values, specifically focusing on angiogenesis. Since in our model, tumor-vessels development depends on AFs distribution, we mainly considered the parameters impacting *af*. Moreover, since we assume *af* at equilibrium at the first simulation step, it was sufficient to compute one single simulation step to determine if vascularization occurred or not.

Among the parameters, we selected those known with higher uncertainty, which are *D*_*a*f_, *V*_*pT*_, and *V*_*uc*_ (see SI). Moreover, since RH growth is slow compared to tumor vascularization, we simulated the PFM for different values of *d*_0_. The result is reported in Figure 3.

**Figure 3.**
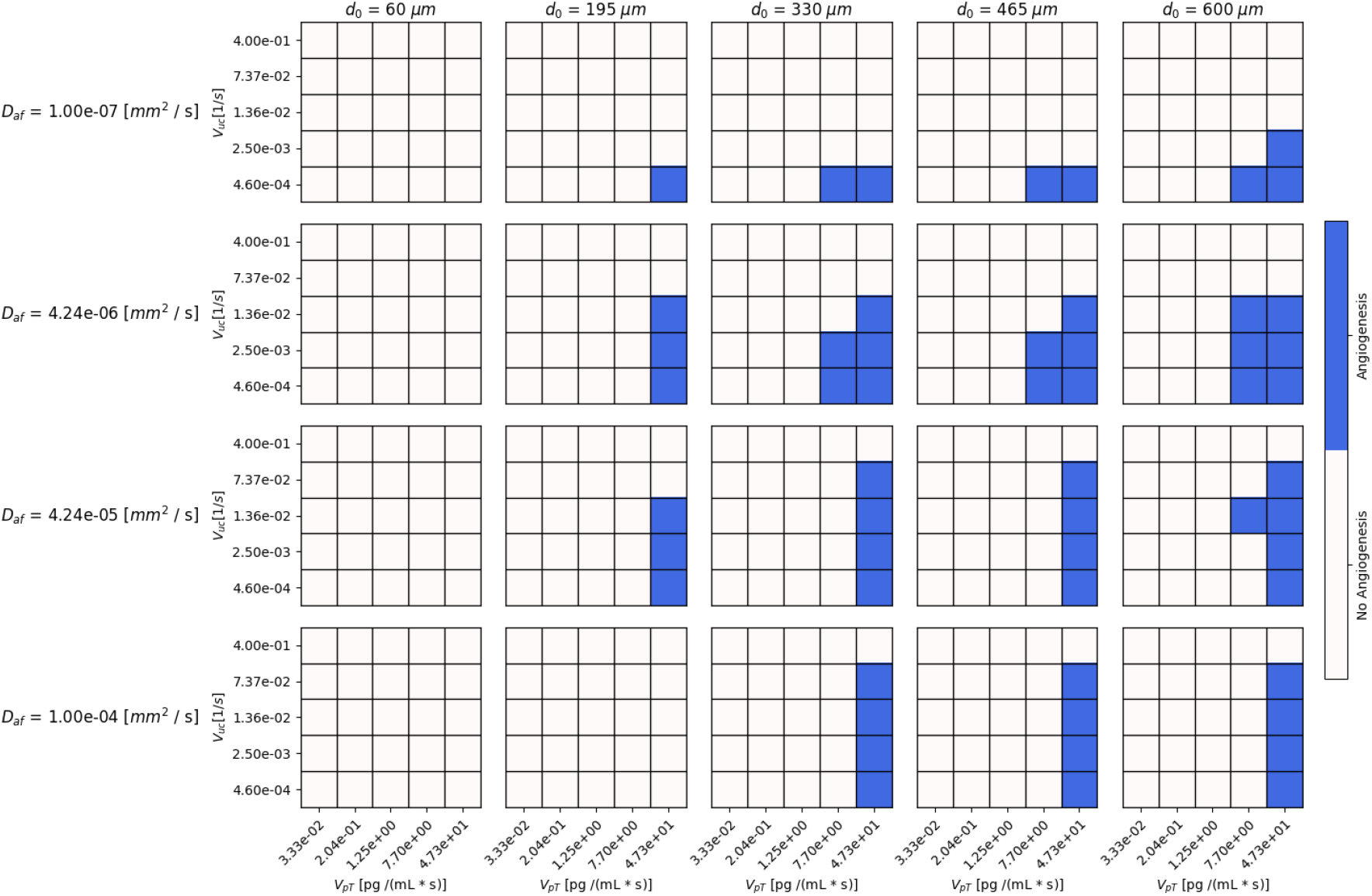
Effect of parameter’s value change on tumor vascularization. We reported a total of 500 simulations divided into 20 grids. Each row of grids represents a set of simulations for a given value of *D*_*af*_; each column for a given value of *d*_0_. Inside each grid, a square represent a simulation for a given value of *V*_*pT*_ (x-axis) and *V*_*uc*_ (y-axis). For the value of the other parameters, see Table 1.

The easiest thing to observe is that a small RH can never induce angiogenesis when its diameter is minimal (*d*_0_ = 1/10 of the RH observed in the patient), regardless AFs’ production or uptake. In any case, AFs concentration is insufficient for such a small tumor to trigger angiogenesis.

For all other *d*_0_ values, novel capillaries are induced only for high AFs production rates (*V*_*pT*_ = 7.7, 47.3 [*pg* · *mL*^−1^ · *s*^−1^]). Since we estimated the tested *V*_*pT*_ values from experimental evidence on non-VHL-related tumor cells (see SI), this result strongly agrees with VHL pathology, characterized by tumor cells overexpressing AFs.

Unsurprisingly, high production rates (*V*_*pT*_) combined with low uptakes (*V*_*uc*_) are the most favorable conditions for tumor-induced angiogenesis in every case. AFs uptake is not critical as the production is for angiogenesis occurrence since we observe TCs’ activation for every *V*_*uc*_ value except the maximum. The reason is that the initial capillaries’ volume is small compared to the RH volume, so differences in orders of magnitude in *V*_*uc*_ do not translate into critical changes for the tumor vascular network.

Less obviously, we observe that diffusivity does seem not to play a critical role in the induction of novel capillaries, at least in the simulated range. However, we observe that both high and low *D*_*af*_ values disfavor angiogenesis. The reason is that in the angiogenesis model we employ there are minimum threshold values for both G and AFs concentration that trigger TCs (see SI and Table 1). On the one hand, a high diffusivity reduces G so that it does not reach the minimum value to trigger the activation of TCs. On the other hand, a low diffusivity generates steeper gradients but also reduces AF concentration outside the tumor volume. Thus, there are situations where it is AFs concentration that is not enough to trigger angiogenesis. Indeed, we observe that *V*_*uc*_ plays a more critical role when D is small.

### RH vascularization might occur very quickly

We already mentioned that angiogenesis occurs rapidly in the simulation reported in Figure 2. Thus, we wondered if the same could happen for different parameter choices. To this end, we selected the parameter sets triggering angiogenesis from Figure 3 and extended the simulations for 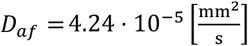 and *d*_0_ = 600 [μ*m*] to observe RH-driven angiogenesis over time. We picked that *D*_*af*_ value because it derives from experimental evidence (see SI), and thus we consider that as the most reliable in the range. The reason for the *d*_0_ value is that we assume that the larger the tumor, the longer the time required for capillaries invasion. For all of them, we simulated 300 time steps (150 h) (Figure 4).

**Figure 4.**
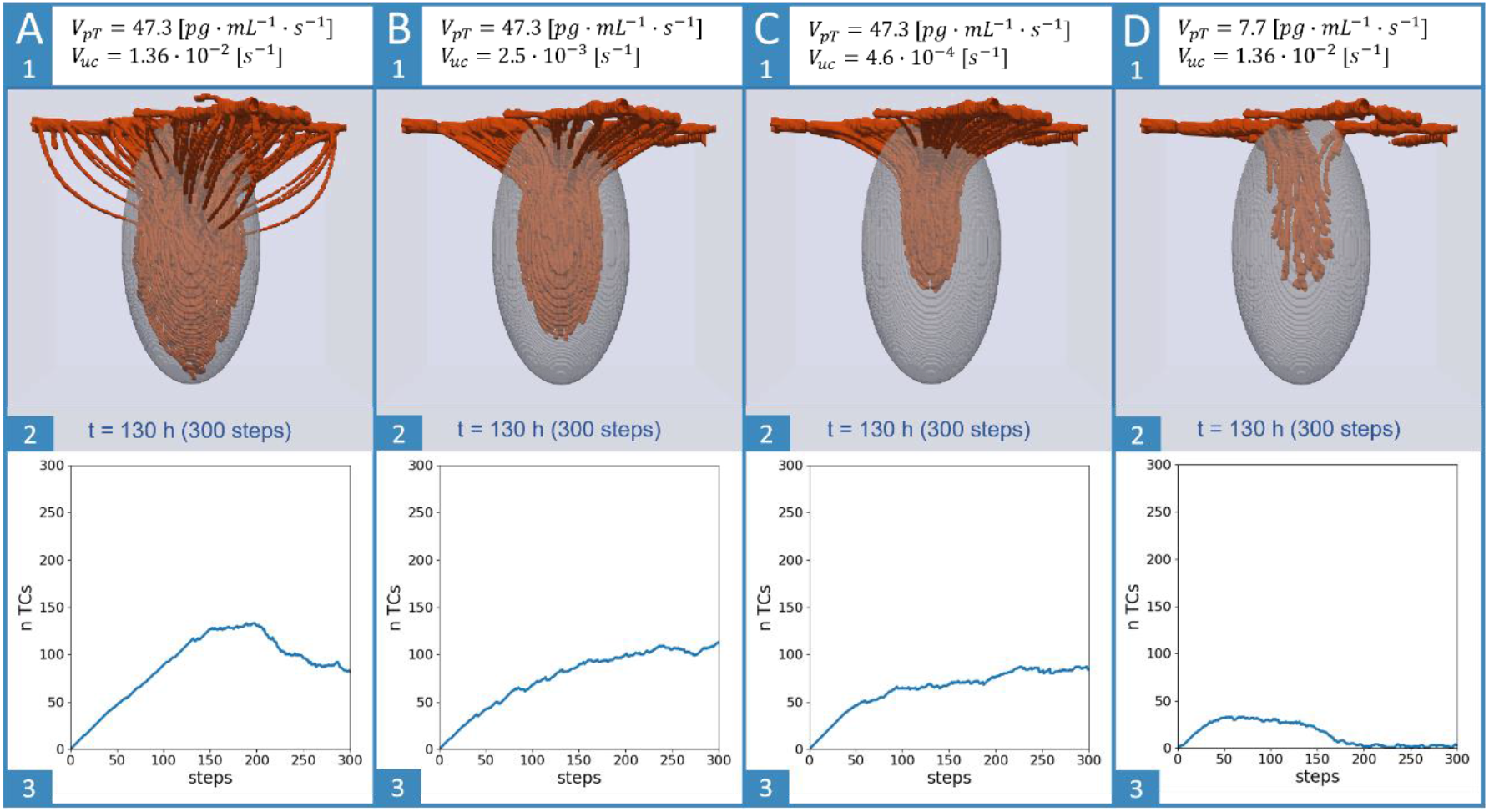
RH vascularization for different parameters set. Four simulations are shown (A, B, C, and D). For each simulation, *d*_0_ = 600[μ*m*] and *D*_*af*_ = 4.24 · 10^−5^[*mm*^2^ · *s*^−1^]. *V*_*pT*_ and *V*_*uc*_ values are shown at the top of each column (A1, B1, C1, and D1). Below (A2, B2, C2, and D2), we reported the capillaries network after 150 h of simulated time. In the last row (A3, B3, C3, and D3) we reported the number of TCs over time.

We observe that vascularization occurs quite rapidly, and sometimes 150 h is enough for complete tumor invasion. Unexpectedly, lower uptake (*V*_*uc*_) disfavors angiogenesis, reducing the number of TCs and their velocity. The reason is that a lower uptake reduces G, which determines both TCs activation and their speed (see Materials and Methods).

A lower production rate (*V*_*pT*_) reduces tumor vascularization, resulting in a sparser capillary network paired with a relatively low vascularization time. Moreover, we observe a lower number of TCs over time. Interestingly, even though full tumor invasion is not reached in 300 steps, the number of active TCs drops after 175 steps, suggesting that the capillary network is already close to stabilization.

## DISCUSSION

In this work, we presented a PFM to simulate RH growth and angiogenesis in a patient-specific way. Despite the model is, as every CMM, a strong simplification of reality, we showed that it is capable to reproduce RH development in time, closely matching with the OCTA images reported for the patient (Figure 2). Moreover, we used our model to show the crucial role that time plays in RH development, which, in the absence of an animal model (Park & Chan, 2012), is challenging to observe otherwise.

First, we note that tumor vascularization does not occur for every tumor diameter, suggesting that RH must reach a minimal dimension to trigger sprouting angiogenesis (Figure 3). Due to the uncertainty associated with the PFM’s parameters we can’t use our model to estimate such dimension, which is also likely to be patient specific. However, such minimal dimension probably exists, since both a minimal AFs concentration and a minimal gradient are necessary to trigger angiogenesis (Travasso, Poiré, et al., 2011). This fact, together with the slow growth rates associated with VHL-related hemangioblastomas (Ye et al., 2012), might explain why we could not produce a murine model for RH. It might be that the short life span of rodents reduces the probability to reach such minimal dimension.

Second, we observe that angiogenesis is triggered only for high AF production rates. Considering that the range of simulated *V*_*uc*_ values derives from measures and estimations in non-VHL-related tumor cells (see SI and (Finley et al., 2013)), we interpret this result as a further confirmation of the validity of our model. Indeed, RH is strongly associated with VHL, and VHL tumors overexpress AFs because their oxygen sensing pathway is impaired. The same might occur in neoplasms also outside the syndrome (Kim & Kaelin, 2004) but, in general, tumor vascularization is triggered under hypoxic condition (Xu et al., 2016). Thus, it makes sense that the AF expression values estimated in non-VHL-tumor cells, with a functioning oxygen sensing pathway, is not always enough to trigger angiogenesis in our case study.

Third, we observed that when angiogenesis is triggered, it takes place rapidly, sometimes invading the full tumor mass in a few days. Even though our simulations consider a patient-specific case, it must be noted that RHs are often very small (1.5 mm or smaller (Karimi et al., 2020)) and that TCs’ velocity has been reported to be up to 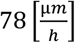 (Blinder et al., 2015).

This fact provides a new perspective on the disappointing results of AAT for this tumor. Despite an initial hope in the effect of this approach to treat early-stage RHs, the clinical evidence reported so far did not match the expectations. Angiogenesis inhibitors have only led to exudation reduction and minimal or absent tumor regression (Wiley et al., 2019). Since the primary purpose of this therapy is the reduction of VEGF concentration in the tumor, it has been proposed that targeting only this molecule might not be enough to prevent vascularization (Wiley et al., 2019). Indeed, other AFs play a relevant role in RH-induced angiogenesis. However, our simulations suggest that time also has a critical role in the effectiveness of AAT. If RH-induced angiogenesis is quick, as predicted by our model, the capillaries may already be too mature to be efficiently targeted with this therapy.

It is already well known that angiogenesis inhibitors mainly induce regression for immature vessels. Indeed, it is commonly used in combination with chemotherapy to stabilize blood vessels and improve the delivery of other drugs. (Vasudev & Reynolds, 2014) Moreover, several facts agree with our suggestion that the therapy might be too late. First, RH is more easily diagnosed only when vascularization is already present. Second, a recent clinical study, which introduced the inhibition of other AFs, has not improved the effect of AAT (Hwang et al., 2021). Finally, a recent case report employed OCTA to display the effect of this therapy on the capillaries of a large RH, showing that larger blood vessels remained stable (Otero-Marquez et al., 2022).

In conclusion, our model is remarkably capable of recapitulating the critical aspects of RH pathology and development in time, which are otherwise very hard to observe in the absence of an animal model. These results suggest that this model could be used to predict RH development and the effect of therapy in a patient-specific way. However, further validation with more OCTA images is necessary to assess the predictive capability of the PFM thoroughly. Our work provides several key insights into RH pathology and the effect of therapy. Moreover, it shows the value of OCTA images to investigate RH (or other vascular pathologies) not only statically but dynamically with the help of mathematical models.

## MATERIALS AND METHODS

### OCTA case report selection

Despite the low incidence of VHL, we could find several case reports (CRs) providing OCTA images of RH. Most are focused on the effect of therapy (Chou et al., 2018; Otero-Marquez et al., 2022; Russell et al., 2020), but some also present treatment-naïve RHs (Custo Greig & Duker, 2021; Goswami et al., 2021; Sagar et al., 2018), allowing direct observation of the angiogenesis induced by the tumor. Among these reports, the one presented by Goswami A. and collaborators (Goswami et al., 2021) was the only one showing evidence of an early-stage RH where both the capillaries and the borders were clearly visible. Moreover, the reported RH was exophilic, with a marked capillary invasion in toward the outer retina. Since this part of the retina is not vascular, this aspect was crucial for distinguishing the capillaries induced by the tumor and the original capillaries in the inner retina. Moreover, inner capillaries showed no sign of enlargement of tortuosity. Thus, we selected the CR presented by Goswami A. *et al*., and we used the OCTA images presented to a) estimate the initial capillary state with relative confidence; and b) validate our model results with patient-specific data.

### Initial capillary network

To construct an image of the putative initial capillaries network (PICN) before tumor-induced angiogenesis, we used GNU Image Manipulation Program (GIMP) to modify the original image reported in the SCR. More precisely, we added a dark layer on the superficial OCTA image to obscure the tumor’s capillaries, of which the prosecution could be seen in the outer retina OCTA image. We kept the feeding arterioles and venules as they showed no sign of enlargement or tortuosity. Of course, we limited our modifications to the proximity of RH (see SI).

### Mesh construction

Our simulations employed a standard box-shaped mesh of tetrahedral elements constructed using the FEniCS computational platform (Alnæs et al., 2015; Logg et al., 2012).

To contain the lesion area, we set the x and y sides of the mesh equal to ∼ 1.4 *mm* (see the red square in Figure 2b and 2c) and the z side equal to 1.2 mm (twice the tumor’s diameter). We set the maximum side length for all tetrahedral elements to 0.7 · *R*_*c*_ to allow the observation of TCs, which are represented as rounded cells of radius *R*_*c*_.

### Blood vessels 3D reconstruction

The procedure we used to 3D-reconstruct the PICN is composed of two main steps. First, we extracted a 2D image of the PICN (2D-PICN); second, we used this image to construct a 3D phase-field of the PICN (3D-PICN) to employ as the initial condition for c.

To obtain the 2D-PICN, we used the Trainable Weka Segmentation (TWS) tool implemented in Fiji (Schindelin et al., 2012). We trained the TWS with two classes, one representing the capillaries (C1) and the other representing the retinal tissue (C2). After the training, we obtained a probability map (PM) associating each pixel with the probability of being in C1. Then, we processed the PM with Scikit-image (Van Der Walt et al., 2014), keeping only the pixels with more than a 50% probability of being C1 and removing isolated pixels. Finally, the image was binarized to obtain the 2D-PICN (see SI for details).

After that, we used 2D-PICN to compute the 3D-PICN. To do so, we developed our own simple and lightweight reconstruction algorithm (RA), which assumes that the local radius of the vessels can be estimated using the local distance between the 2D vessel axis and its edge. We refer to the SI for a detailed discussion of the RA.

### Mathematical model for angiogenesis

To simulate tumor vascularization, we employed a mathematical model already presented in the literature (Travasso, Poiré, et al., 2011), which was slightly adapted to better represent our case study. In this section we briefly present the basic concepts behind this model and the small variations we introduced.

The approach proposed by Travasso and collaborators merges a PFM with an agent-based algorithm, which rules the tip cells (TCs) and sprouting angiogenesis. The PFM component is a PDE defining the evolution in time of *c*:

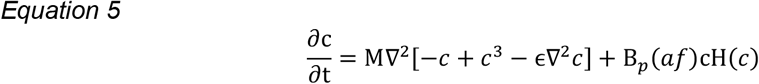

Where the first term is a Cahn-Hillard term accounting for the interface dynamics, and the second term accounts for endothelial cells’ proliferation. AFs regulate the latter through the B_p_(af) function, which is defined as:

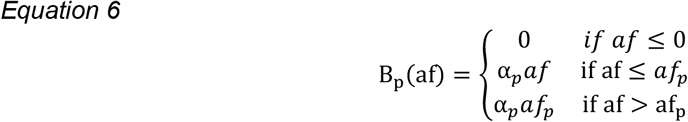

The agent-based algorithm handles TCs’ activation, deactivation, motion, and stalk cells’ (SC) proliferation. We refer to the original publication for an extensive explanation of the algorithm, but we also reported some details in SI.

To merge the agents with the PDE, at each time step, any point of *c* which is inside a TC (i.e., closer than *R*_*c*_ to the position of a TC agent) is updated to the following value:

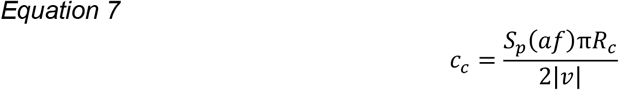

Which accounts for the TC mass and the SCs’ proliferation. The latter is coupled with AFs with the function *S*_*p*_(*af*), which has the same definition of *B*_*p*_(*af*) but uses different parameters:

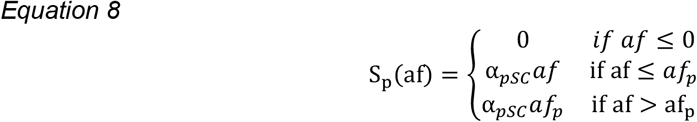

The original model did not include this distinction, and we added it to differentiate the SCs’ proliferation from mature endothelial cells’ proliferation (see SI for details).

As described above, we used the PICN as the initial condition. We employed natural Neumann conditions at the boundaries:

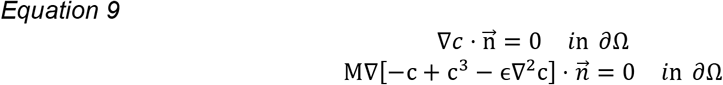

The use of such boundary conditions implies that any local variation of *c* is due to proliferation and sprouting angiogenesis, without any transport through the mesh surface.

### Numerical Methods and Implementation

We ran our simulation exploiting FEniCS, a computational platform to write and simulate PDE-based models (Alnæs et al., 2015; Logg et al., 2012). To include angiogenesis in our model, we exploited the open implementation in the Python package Mocafe (Pradelli et al., 2022).

To solve our PDE system, we employed a standard spatial discretization using Lagrange finite elements of the first order. These elements are C^1^, so they cannot handle fourth-order PDE like the one used for capillaries. Thus, we separated this equation into two second-order PDEs, introducing an auxiliary variable μ:

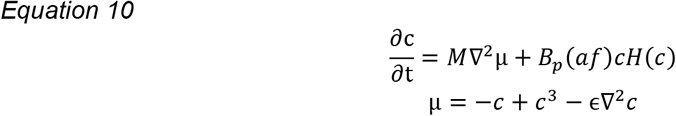

For temporal integration, we used a standard implicit Euler scheme. Our time step was constant and equal to about 26 min (see Table 1).

Since our PDE system is non-linear, we used the Newton-Raphson method to linearize the algebraic system resulting from the spatial and temporal discretization. Then, we used the generalized minimal residual method (GMRES) (Saad & Schultz, 2006) with an algebraic multigrid preconditioner (AMG) to solve the system at each time step.

### Visualization

To visualize our simulations’ result, we used ParaView (Ahrens et al., 2005). More precisely, we always employed the isosurface *c* = 0 to display the capillaries and the isosurface φ = 0.5 to display the tumor. The simulated RH sections shown in Figure 2g and 2i are obtained using the Clip filter in ParaView. Finally, we created the TCs plot in Figure 4 and the tiles plot in Figure 3 using the Matplotlib Python package (Hunter, 2007).

## ACKNOWLEDGMENTS

The authors thank Associazione Italiana per la Ricerca sul Cancro (grant nr. IG 2019 ID. 23825) for funding this research project. We also thank Prof. Dr. Pradeep Venkatesh for helpful insights regarding the selected case report.

## DATA AVAILABILITY STATEMENT

The code used in this manuscript is freely available under a CC BY-NC 4.0 license on GitHub (https://github.com/fpradelli94/rh_mocafe) and Zenodo (https://doi.org/10.5281/zenodo.7330072). The authors commit to the long-term maintenance of the code.

