## Supplementary Information for "Patient-specific simulation of Retinal Hemangioblastoma provides new perspectives on the role of antiangiogenic therapy"

**Other supporting materials:**

- Movie S1

### SECTION 1: MATHEMATICAL MODEL

**Detailed mathematical model.** Our mathematical model is composed of the following equations:

| | |
|---|---|
| Retinal Hemangioblastoma (RH) Growth | *Equation s1* $$\varphi(\vec{\boldsymbol{x}},\mathrm{t}) = \begin{cases} 1 & \text{if } \left(\frac{\mathrm{x}}{\mathrm{s_x(t)}}\right)^2 + \left(\frac{\mathrm{y}}{\mathrm{s_y(t)}}\right)^2 + \left(\frac{\mathrm{z}}{\mathrm{s_z(t)}}\right)^2 \leq 1 \\ 0 & \text{Otherwise} \end{cases}$$ Where: *Equation s2* $$\mathrm{s_x} = \mathrm{s_y} = \frac{\mathrm{d_0}}{2}\mathrm{tg}r^{\frac{\mathrm{t}}{3}} \qquad s_z = d_0\mathrm{tg}r^{\frac{\mathrm{t}}{3}}$$ |
| Capillaries PDE | *Equation s3* $$\frac{\partial \mathrm{c}}{\partial \mathrm{t}} = \mathrm{M}\nabla^2[-c + c^3 - \epsilon\nabla^2 c] + \mathrm{B}_p(af)\mathrm{cH}(c)$$ Where: *Equation s4* $$\mathrm{B_p(af)} = \begin{cases} 0 & if\ af \leq 0 \\ \alpha_p af & \text{if af} \leq af_p \\ \alpha_p af_p & \text{if af} > \mathrm{af_p} \end{cases}$$ |
| Tip Cells (TC) velocity | *Equation s5* $$\mathrm{v} = \begin{cases} \chi\frac{\nabla \mathrm{af}}{\mathrm{G}} & \text{if } \mathrm{G_m} \leq \mathrm{G} < \mathrm{G_M} \\ \chi\frac{\nabla \mathrm{af}}{\mathrm{G}}\mathrm{G_M} & \text{if } \mathrm{G} \geq \mathrm{G_M} \end{cases}$$ |
| Tip Cells (TCs) and Stalk Cells (SCs) value | *Equation s6* $$c_c = \frac{S_p(af)\pi R_c}{2\lvert v \rvert}$$ Where: *Equation s7* $$\mathrm{S_p(af)} = \begin{cases} 0 & if\ af \leq 0 \\ \alpha_{pSC} af & \text{if af} \leq af_p \\ \alpha_{pSC} af_p & \text{if af} > \mathrm{af_p} \end{cases}$$ |

| Angiogenic Factors (AFs) | *Equation s8* $$\frac{\partial af}{\partial t} = D_{af}\nabla^2 af + V_{pT} \cdot \varphi \cdot \left(1 - H(c)\right) - V_{uc} \cdot af \cdot H(c) - V_d \cdot af$$ |
|---|---|

Plus, there are algorithms taking care of tip cells' (TCs) activation and deactivation, which we represented in Figure S6.

### SECTION 2: PARAMETERS

Each of the equations and algorithms presented above depend on several parameters, which, when possible, were based on experimental evidence. In the following, we describe how we derived each parameter. Note that we often estimated the parameters regarding the angiogenic factors (AFs) using the vascular endothelial growth factor (VEGF) as the primary reference. Indeed, the association of this cytokine with tumor-induced angiogenesis is widely recognized (Kut et al., 2007); moreover, there is experimental evidence on the fact that VEGF is overexpressed in RH (Los et al., 1997).

On the GitHub repository related to this manuscript, you can also find a Jupyter Notebook (https://github.com/fpradelli94/rh_mocafe/blob/main/notebooks/parameters.ipynb) to make the evaluation of each parameter fully reproducible.

In the following, we will refer to the case report presented by Goswami et al. (2021) as the selected case report (SCR) for brevity.

**Arbitrary Units.** We choose 800 μm as space arbitrary unit (sau), which is of the same order of magnitude of RH diameter at its maximum extent (see section $\boldsymbol{d_0}$ **(Initial RH diameter)**).

As time arbitrary unit (tau) we assumed 26 min, which is the time step value used to solve the PDE system. The same value is used by Travasso and collaborators in the original model (Travasso et al., 2011) and allows the integration between the PFM model and the algorithm for TC. Indeed, given that the maximum TCs' velocity (see sections $\boldsymbol{G_m}$, $\boldsymbol{G_M}$, and $\boldsymbol{\chi}$ (Tip cells velocity)) is:

$$v_{max} = 0.35 \left[\frac{\mu m}{min}\right]$$

The maximum displacement of a TC in one time step is:

$$s_{max} = v_{max} \cdot dt = 9.1\ [\mu m]$$

Which is less than one TC radius (see section $\boldsymbol{R_c}$ (Tip Cells' radius)). In this implementation, using a higher value of dt could lead to a discontinuous $c$ field.

For AFs concentration, we assumed an arbitrary unit equal to 6 $\left[\frac{ng}{mL}\right]$. This value represents a relatively high concentration value. It was selected based on the maximum VEGF concentration reported by Na X. and collaborators (Na et al., 2003), which measured VEGF concentration in some VHL-related kidney tumors. Thus, it looked like a reasonable measure unit for a VHL-related tumor like RH.

$\boldsymbol{d_0}$ **(Initial RH diameter).** The SCR clearly shows the lesion dimension but does not provide precise spatial information. Thus, we contacted Prof. Dr. Venkatesh (the corresponding author for the SCR), who kindly provided us with the original images of the lesion. It is common to use the patient's optical disk as a reference to estimate the dimension of RHs, and we did the same using the images provided by the original author. We overlapped the Optical Coherence Tomography Angiography (OCTA) with the image showing the lesion with OD. We estimated the lesion diameter to be about 1/3 of the vertical axis (see Figure S7). Given that such an axis is typically 1.88 mm long (Arora et al., 2015), we estimated the RH diameter to be 0.6 mm (600 μm).

In agreement with this estimate, the initial tumor's diameter must be lower. Thus, we performed our simulations with $d_0$values ranging from 60 to 600 μm.

**$tgr$ (Tumour growth rate).** The average growth rate for VHL-related hemangioblastoma has been reported to be 0.35% per year (considering the tumor volume's increase) (Ye *et al.*, 2012). Therefore, we employed this value for our model.

**M (Motility for c field).** This value represents the motility of the capillaries phase field $c$, which we assumed to be in line with the value used by Travasso et al. (Travasso et al., 2011) in the original paper ($\mathrm{M} = 10^{15}\ \frac{\mathrm{m}^2}{\mathrm{s}}$).

**$\epsilon$ (Interface width for the c field).** This value represents a measure of the stable interface's width for the capillaries phase field c, which we assumed to be equal to the value used by Travasso et al. (Travasso et al., 2011) in the original paper ($\epsilon = 1.25^2 \mu m^2$).

**$B_p$ (Proliferation for mature endothelial cells) and $S_p$ (proliferation for stalk cells).** In the original model (Travasso et al., 2011), there was no distinction between mature endothelial cells' and stalk cells' proliferation. They considered only proliferation due to stalk cells (SCs), for which they assumed:

$$\alpha_p \cdot af_p = 0.97\ \left[\frac{1}{hr}\right] \qquad \mathrm{af}_p = 0.3\ [afau] \qquad \alpha_p = \frac{0.97}{0.3}\ \left[\frac{1}{hr \cdot afau}\right]$$

Which represents the maximum proliferation for the SCs. Notice that this value represents an increase in volume per unit time and not an exact proliferation rate for SCs; however, its choice is based on *in vitro* evidence, as justified by Travasso and collaborators (Travasso et al., 2011). Thus, we kept the same parameters values for SCs' proliferation:

$$\alpha_{pSC} \cdot af_p = 0.97\ \left[\frac{1}{hr}\right] \qquad \mathrm{af}_p = 0.3\ [afau] \qquad \alpha_{pSC} = \frac{0.97}{0.3}\ \left[\frac{1}{hr \cdot afau}\right]$$

However, for mature endothelial cells, we neglected the proliferation induced by AFs, assuming:

$$\alpha_p = 0$$

This choice has two reasons. First, performing simulations with several $\alpha_\mathrm{p}$ values (e.g., $= \alpha_{pSC}$, $= \frac{\alpha_{pSC}}{10}$), we observed capillaries enlargement instead of sprouting angiogenesis, demonstrating that our model was not correctly reproducing the SCR for these parameter choices (see Figure S8). Second, mature blood vessels are more stable than novel capillaries, and the proliferation of mature endothelial cells follows different rules, which we choose not to cover in our model (Vasudev & Reynolds, 2014). Thus, it is reasonable to assume that mature endothelial cells' proliferation is negligible compared with SCs' proliferation.

**$G_m$, $G_M$, and $\chi$ (Tip cells velocity).** Equation s5 defines the TCs' velocity range, which has been measured both *in vitro* and *in vivo*. Travasso and collaborators (Travasso et al., 2011) assumed the maximum value for TCs' velocity to be:

$$v_{max} = 0.35\ \left[\frac{um}{min}\right]$$

This value was derived from experimental evidence *in vitro* (Stokes et al., 1991). Moreover, it agrees with the TCs' velocity measured during sprouting angiogenesis (Blinder et al., 2015), and *in vivo* experiments (Guedez et al., 2003; Harper et al., 2021).

$G_m$ represents the minimum value of G necessary to activate the tip cells. The lowest VEGF gradient triggering endothelial cells' migration we could find in the literature is $14\ \left[\frac{ng}{mL \cdot mm}\right]$ (Shamloo et al., 2008). Thus, we assumed:

$$G_m = 14\ \left[\frac{ng}{mL \cdot mm}\right]$$

Then, coherently with the original model, we assumed $G_M = 3G_m$:

$$G_M = 42\ \left[\frac{ng}{mL \cdot mm}\right]$$

This leads to the assumption that TCs' velocity linearly increases with G in the range [14, 42] ng / (mL mm). The fact that TCs' velocity depends on G over a limited range has been proved experimentally by Barkefors I. and collaborators (Barkefors et al., 2008). They also provide approximate G values necessary for endothelial cells' migration, which are slightly higher but in agreement with ours (about $[50,\ 100]\,\frac{ng}{mL \cdot mm}$).

Finally, we assume that χ is such to limit the maximum TCs' velocity to $v_{max}$, so:

$$\chi = \frac{v_{max}}{G_M}$$

$\boldsymbol{R_c}$ **(Tip Cells' radius).** Assuming the maximum value provided on the BioNumbers database (Entry ID: 100432; (Milo et al., 2010)), we have:

$$R_c = 10\ [\mu m]$$

$\boldsymbol{T_c}$ **(Minimum AFs concentration for tip cells activation).** Despite being difficult to measure, a threshold in AFs concentration for TCs' activation must exist since this phenomenon is triggered by AFs binding to endothelial cells' surface receptors. Here, we assumed that AFs concentration must be in the same order of magnitude as the local concentration of the receptors to generate TCs activation.

Imoukhuede and collaborators have measured the number of VEGF receptors (VEGFRs) per cell in different conditions, finding that their number is in the range: $[10^3, 10^4]\left[\frac{VEGFR\ molecules}{cell}\right]$ (Imoukhuede & Popel, 2011). We can translate this quantity in a spatial concentration estimating the number of endothelial cells per unit volume, i.e., the cell density ($CD$).

Assuming each cell to have about the same volume as a sphere with radius $R_c$, we can assume a cell volume to be around $\frac{4}{3}\pi R_c^3$. Thus, if we consider a portion of space filled with endothelial cells (e.g., the capillary wall), the number of cells per unit volume is:

$$CD_{max} = \frac{1}{\frac{4}{3}\pi R_c^3} \cong 2.4 \cdot 10^{-4}\left[\frac{cells}{\mu m^3}\right] \cong 2.4 \cdot 10^{8}\left[\frac{cells}{mL}\right]$$

If we want to be more conservative, we can assume that only 75% of space in the capillary wall is composed of cells, while the rest is the extracellular matrix. This leads to a slightly lower estimation:

$$CD_{min} = \frac{0.75}{\frac{4}{3}\pi R_c^3} \cong 1.8 \cdot 10^{-4} \left[\frac{cells}{\mu m^3}\right] \cong 1.8 \cdot 10^{8} \left[\frac{cells}{mL}\right]$$

With this estimate, we can estimate the range of VEGFRs concentration in the capillary wall:

$$VEGFR_{max} = \frac{10^4 \cdot CD_{max}}{c_{Avogadro}} \cong 4 \cdot 10^{-12} \left[\frac{moles}{mL}\right]$$

$$VEGFR_{min} = \frac{10^3 \cdot CD_{min}}{c_{Avogadro}} \cong 3 \cdot 10^{-13} \left[\frac{moles}{mL}\right]$$

And we assume that AFs concentration must be of the same order of magnitude to activate TCs:

$$T_c \in [3 \cdot 10^{-13}, 4 \cdot 10^{-12}] \left[\frac{moles}{mL}\right]$$

Considering that $T_c$ represents a minimum threshold value, we assume that it is enough to have it in the order of $10^{-13} \left[\frac{moles}{mL}\right]$.

We can translate this value to $\frac{pg}{mL}$ by knowing the molecular weight of VEGF. UniProt reports $16\ kDa$ for the smallest VEGF isoform (identifier: P15692-10) and $45\ kDa$ for the biggest (identifier: P15692-14) (Bateman et al., 2021). Thus, considering the average, we have:

$$T_c = 10^{-13} \cdot \frac{16 + 45}{2} \cdot 10^3 \cong 3000 \left[\frac{pg}{mL}\right]$$

$\boldsymbol{\delta_4}$ **(Minimum TCs distance due to the Notch pathway).** Coherently with the original model (Travasso et al., 2011), we assume that the Notch pathway prevents the activation of two neighbor cells. Thus, we assume the minimum TC distance to be equal to $4R_c$:

$$\delta_4 = 4R_c = 40\ [\mu m]$$

$\boldsymbol{D_{af}}$ **(AFs diffusivity).** Even considering VEGF alone, we could not find an agreed diffusivity value among different publications. VEGF diffusivity has been estimated in several publications (see Table S1) with values ranging from $10^{-7}$ to $10^{-4} \left[\frac{mm^2}{s}\right]$.

Thus, we used this entire range as a reference for our simulations. Additionally, we derived our estimation based on the experiments reported by Kihara et al. (Kihara et al., 2013).

In (Kihara et al., 2013), the authors measured the diffusivity of different biomolecules with different molecular weights. Among the different molecules, they tested Alexa488-dextran and FITC-dextran, which have molecular weights of 10 kDa and 40 kDa respectively. Given that the different isoforms of VEGF have a molecular weight included in the range [16, 45] kDa (derived from UniProt (Bateman et al., 2021); identifiers P15692-10 and P15692-14), we assumed VEGF diffusivity to be in the same range.

More precisely, considering the average molecular weight of 30.5 kDa, we can find an estimation of VEGF diffusivity by interpolating the two values for Alexa488-dextran and FITC-dextran:

$$D_{VEGF} \cong \frac{D_{\text{FITC-dextran}} - D_{\text{Alexa488-dextran}}}{40 - 10} \cdot (30.5 - 10) + D_{\text{Alexa488-dextran}}$$

Resulting in:

$$D_{VEGF} = D_{af} = 4.24 \cdot 10^{-5} \left[\frac{mm^2}{s}\right]$$

$\boldsymbol{V_{pT}}$ **(AFs production rate inside the tumor).** Tumor VEGF secretion has been measured *in vitro* and has been estimated *in vivo* by Finley S. and collaborators (Finley et al., 2013). They estimated VEGF secretion *in vivo* to be in the range $[0.007, 0.023] \left[\frac{molecules}{cell \cdot s}\right]$, while *in vitro* measures have evaluated it to be in the range $[0.03, 2.65] \left[\frac{molecules}{cell \cdot s}\right]$.

Since an impaired VEGF regulation often characterizes von Hippel-Lindau-related tumors, we assumed the VEGF production rate to include both the in vitro and the in vivo range, so to have $V_{pT} \in [0.007, 2.65] \left[\frac{molecules}{cell \cdot s}\right]$.

To convert this range to $\left[\frac{pg}{mL \cdot s}\right]$, we considered the VEGF molecular weight as registered on UniProt (Bateman et al., 2021), which reports $16\ kDa$ for the smallest VEGF isoform (identifier: P15692-10) and $45\ kDa$ for the biggest (identifier: P15692-14). Considering this as the weight of one mole of VEGF, we have:

$$V_{pT_{min}} = \frac{16 \cdot 10^3}{c_{Avogadro}} \cdot 0.007 = 0.02 \cdot 10^{-20} \left[\frac{g}{cell \cdot s}\right]$$

$$V_{pT_{max}} = \frac{45 \cdot 10^3}{c_{Avogadro}} \cdot 2.65 = 19.8 \cdot 10^{-20} \left[\frac{g}{cell \cdot s}\right]$$

Then, we estimated the cell density per unit volume (CD). Assuming a tumor cell to have the same volume as a sphere of radius $R_c$, we have that one volume unit full of tumor cells contains:

$$CD_{max} = \frac{1}{\frac{4}{3}\pi R_c^3} \cong 2.4 \cdot 10^{-4} \left[\frac{cells}{\mu m^3}\right] \cong 2.4 \cdot 10^8 \left[\frac{cells}{mL}\right]$$

If we want to be more conservative, we can assume that only 75% of space in the capillary wall is composed of cells, while the rest is the extracellular matrix. This leads to a slightly lower estimation:

$$CD_{min} = \frac{0.75}{\frac{4}{3}\pi R_c^3} \cong 1.8 \cdot 10^{-4} \left[\frac{cells}{\mu m^3}\right] \cong 1.8 \cdot 10^8 \left[\frac{cells}{mL}\right]$$

Thus, we have:

$$V_{pT_{min}} = 1.8 \cdot 10^8 \cdot 0.02 \cdot 10^{-20} = 0.036 \left[\frac{pg}{mL \cdot s}\right]$$

$$V_{pT_{max}} = 2.4 \cdot 10^8 \cdot 19.8 \cdot 10^{-20} = 47.5 \left[\frac{pg}{mL \cdot s}\right]$$

Finally, the conversion to $\left[\frac{afau}{tau}\right]$ leads to the range:

$$V_{pT} \in [0.0087, 12.3] \left[\frac{afau}{tau}\right]$$

$\boldsymbol{V_{uc}}$ **(AFs uptake by the capillaries).** The AFs uptake factor represents the sum of the biophysical phenomena which result in AFs draining from the lesion. The blood flow plays a critical role in AFs transport, but other factors, such as AFs' receptors uptake and platelets uptake, cannot be neglected at such a small scale. Thus, it is hard to find an experimental reference for the value (or range) of $V_{uc}$.

Thus, we used different reasoning to estimate a range for this parameter. First, we assumed that $V_{uc}$ must be higher than the natural degradation rate for VEGF, $V_d$ (see section $\boldsymbol{V_d}$ (AFs degradation factor)). This is not only realistic, but also necessary for the model to provide a realistic representation of angiogenesis. If $V_{uc}$ is equal to or lower than $V_d$, the AF gradient at the capillary's edges would be zero (if $V_{uc} = V_d$) or directed away from the tumor, and the chemotactic stimulus would be unrealistic. Therefore, we estimate the minimum value for $V_{uc}$ to be double the minimum $V_d$ value reported for VEGF (see section $\boldsymbol{V_d}$ (AFs degradation factor))

$$V_{uc_{min}} = 2 \cdot 0.83 = 1.66 \left[\frac{1}{hr}\right] = 4.6 \cdot 10^{-4} \left[\frac{1}{s}\right]$$

To estimate the maximum value, we selected a value leading to no TCs activation for any combination of the other parameters' values (see Results in the main text). Indeed, the selected case report we aim to reproduce is highly vascular, and TCs activation must have occurred. Thus, any $V_{uc}$ value not triggering sprouting angiogenesis is not interesting for our study.

The selected value was obtained increasing of 100 times the uptake rate in the original model ($0.004 \left[\frac{1}{s}\right]$ (Travasso et al., 2011)):

$$V_{uc_{max}} = 0.4 \left[\frac{1}{s}\right]$$

$\boldsymbol{V_d}$ **(AFs degradation factor).** According to Vempati et al. (Vempati et al., 2014) the VEGF degradation rate is in the range $[0.83, 1.008] \left[\frac{1}{\mathrm{hr}}\right]$. Thus, we assumed the average $0.918 \left[\frac{1}{hr}\right]$ as reference value.

### SECTION 2: METHODS

**Putative initial capillaries network (PICN) estimation.** We constructed the initial capillaries network as explained in the Materials and Methods section in the manuscript. We reported the result in Figure S9. Notice that the modified area (in the centre of Figure S9B) is darker than other areas. However, this cannot influence the result because Figure S9A and Figure S9B are filtered in the following step (see section 2D-PICN extraction).

**2D-PICN extraction.** To extract the PICN from the image reported in Figure S9B, we used the Weka Segmentation implemented in Fiji (Schindelin et al., 2012). More precisely, we used the original image to train the filter. Then we used it to generate a probability map associating the probability of being inside a capillary to each pixel. The result is reported in Figure S10.

After the initial segmentation, we selected the part of the picture corresponding to the simulation area (Figure S11A1). We used Scikit-image (Van Der Walt et al., 2014) to keep only the pixels with more than a 50% probability of being inside a capillary (Figure S11A2). Then, we removed small discontinuities to obtain the binarized reconstruction of the PICN (2D-PICN, Figure S11A3). Finally, we extracted the skeleton and the edges of 2D-PICN to reconstruct the tridimensional PICN in 3D (3D-PICN, Figure S11B). The procedure is reproducible using a Jupyter Notebook (https://github.com/fpradelli94/rh_mocafe/blob/main/notebooks/vessels_image_processing.ipynb) included in the GitHub repository related to this manuscript.

**3D Reconstruction Algorithm (RA).** We depicted a schematic representation of our RA in Figure S12. The output of RA is a Phase-Field, i.e., a 3D scalar field which equals 1 inside the capillaries and -1 outside. Thus, each mesh point must be mapped to 1 or -1 accordingly. For brevity, we will refer to this scalar field as 3D-PICN.

The procedure consists of 3 main steps:

1. *Rescaling 2D-PICN*

   In this step, the 2D-PICN described in section "2D-PICN extraction" is rescaled. The output is a 2D scalar field equal to 1 inside the capillaries and to -1 outside (see Figure S12A).

2. *Setting 2D-PICN to z0*

   Then, we assumed that the 2D-PICN represents a section of 3D-PICN at a given height, z0. Thus, the second step sets the mesh points at height z0 equal to the points of the rescaled 2D-PICN. In our case, we assumed z0 to be 80 μm under the retinal surface because we knew that our initial blood vessels were close to it.

3. *Mapping all other points to 1 or -1.*

   All other mesh points are mapped to 1 or -1 according to the algorithm in Figure S12B. For every given point P, the algorithm works as follows:

   a. It computes the projection of the point P on the plane z0 ($P_{z0}$) and determines its value based on the rescaled 2D-PICN. If the value at $P_{z0}$ is -1, the point P cannot be inside a capillary in 3D. Thus, the algorithm maps P to -1 and stops immediately. Notice that a projection of a point $P = (x_P, y_P, z_P)$ is computed simply by changing $z_P$ to z0, so to have: $P_{z0} = (x_P, y_P, z0)$.

   b. If $P_{z0}$ is 1, the algorithm employs the skeleton and the edges of 2D-PICN to compute the Closest Edge Point (CEP) and the Closest Center Point (CCP). The former is the closest mesh point belonging to the 2D edge of the vessel on 2D-PICN, while the latter is the closest mesh point belonging to the skeleton of 2D-PICN (assumed as the vessel axis).

   c. Once the CEP and the CCP are known, the rule is simple: if the distance between P and CCP is bigger than the distance between CEP and CCP, then P is mapped to -1. Else, P is mapped to 1. This rule allow obtaining approximately cylindrical vessels, with a section close to the local section on 2D-PICN.

A graphical representation of the mapping procedure is reported in Figure S13.

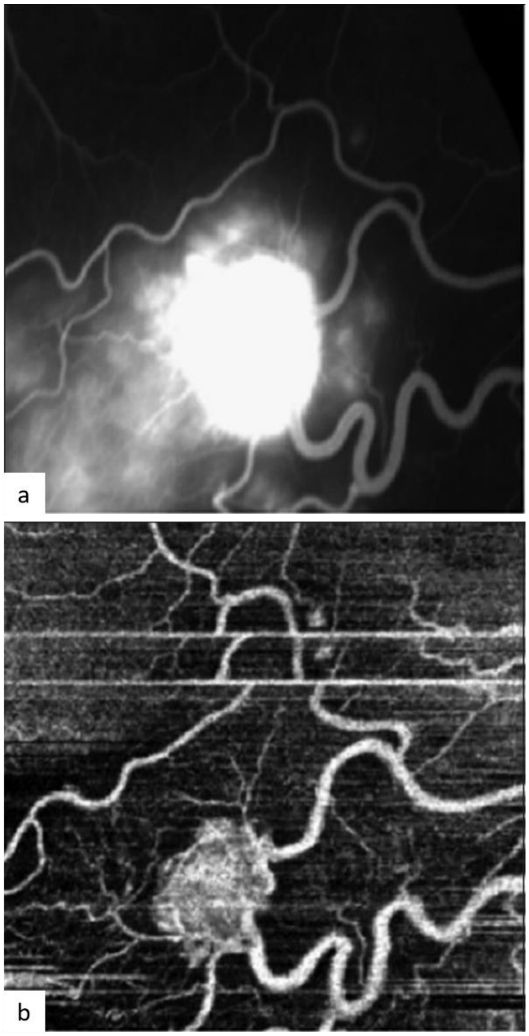


**Fig. S1.** Large RH reported by Sagar P. and collaborators (Sagar et al., 2018). a) Fundus Angiography shows leakage and exudation around the tumor but does not allow a clear observation of the tumor borders and of the capillaries. b) OCTA image displays tumor borders, high vascularity, and major blood vessels enlargement and tortuosity.

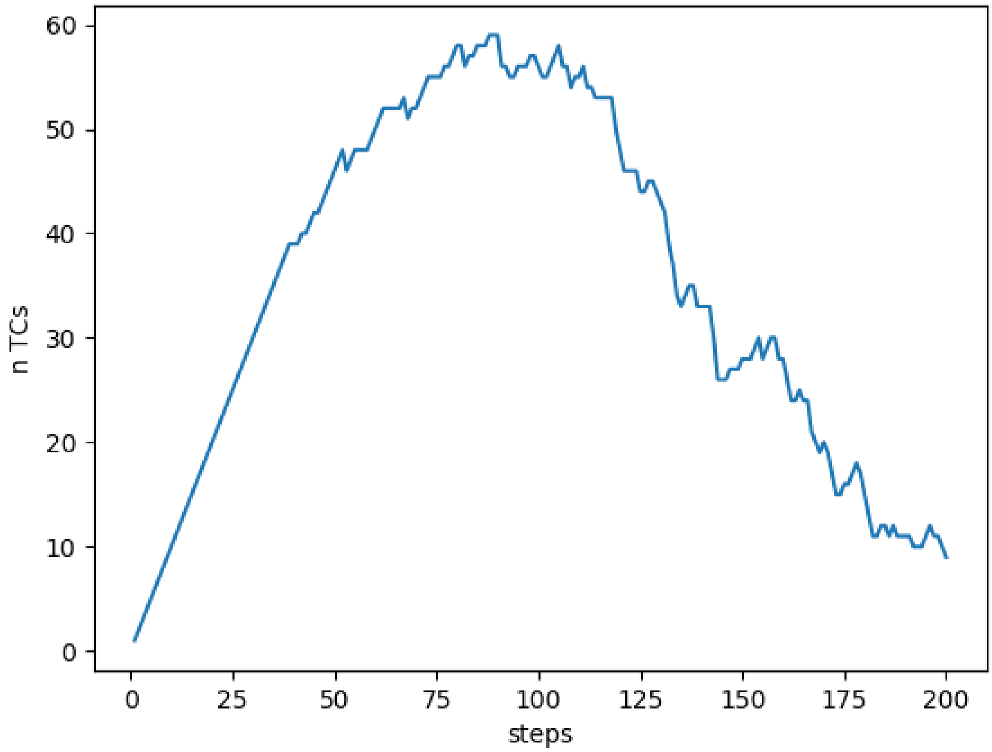


**Fig. S2.** Number of TCs over time for the simulation reported in Figure 2 in the main text.

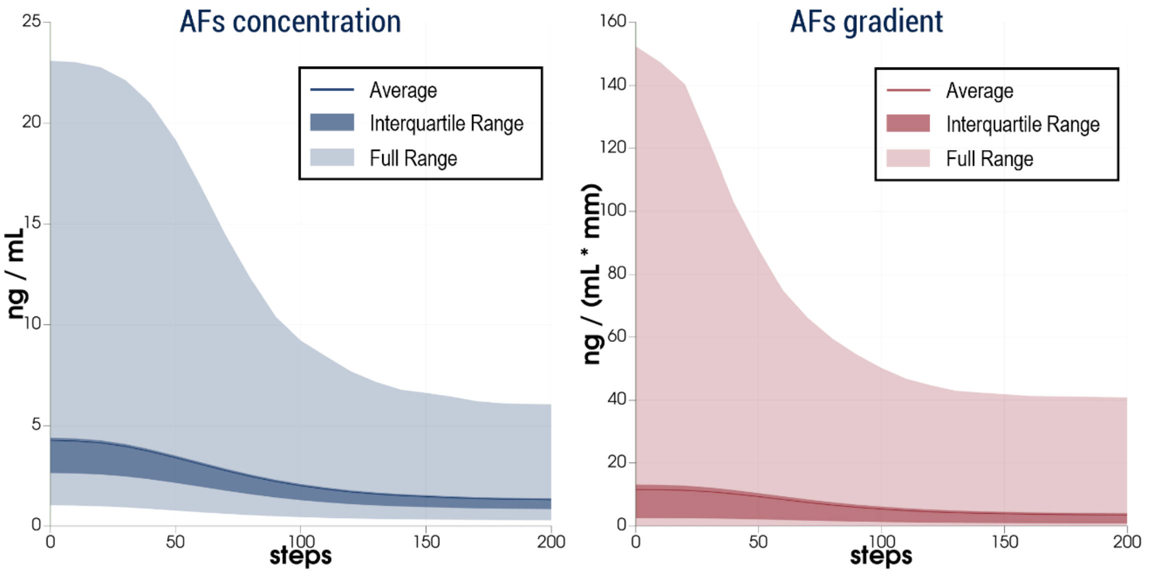


**Fig S3.** AFs concentration and gradient over time for the simulation reported in Figure 2 in the main text.

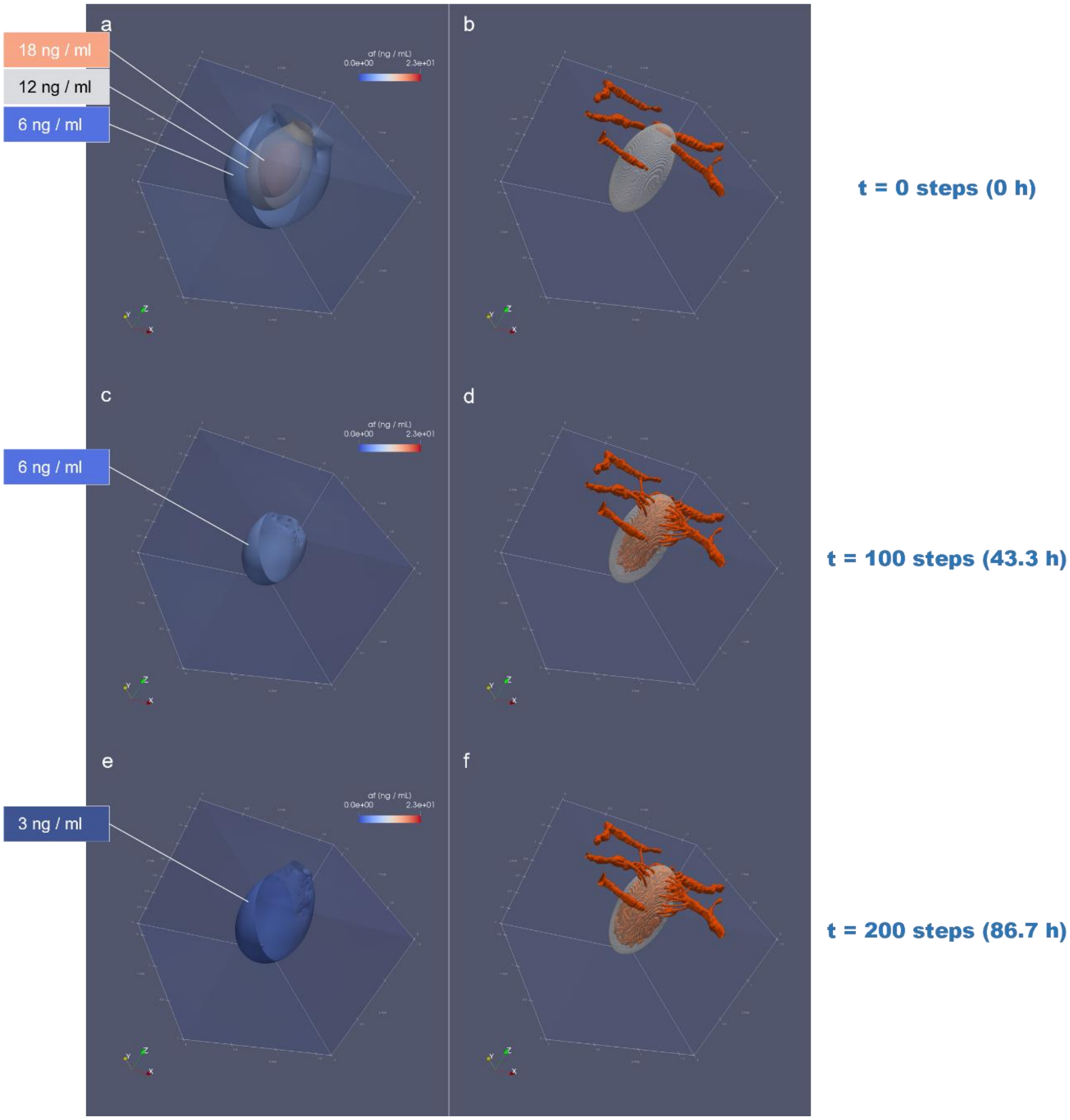


**Fig. S4.** AFs concentration in time (a, c, e) during angiogenesis progression (b, d, f) for the simulation reported in Figure 2 of the main text. In figure a, c, and e, isosurfaces are shown for certain AFs concentration (18, 12, 6, and 3 ng / mL) AFs concentration yelds to 23 ng / mL at the beginning, but in time decreases due to the increasing vascularization.

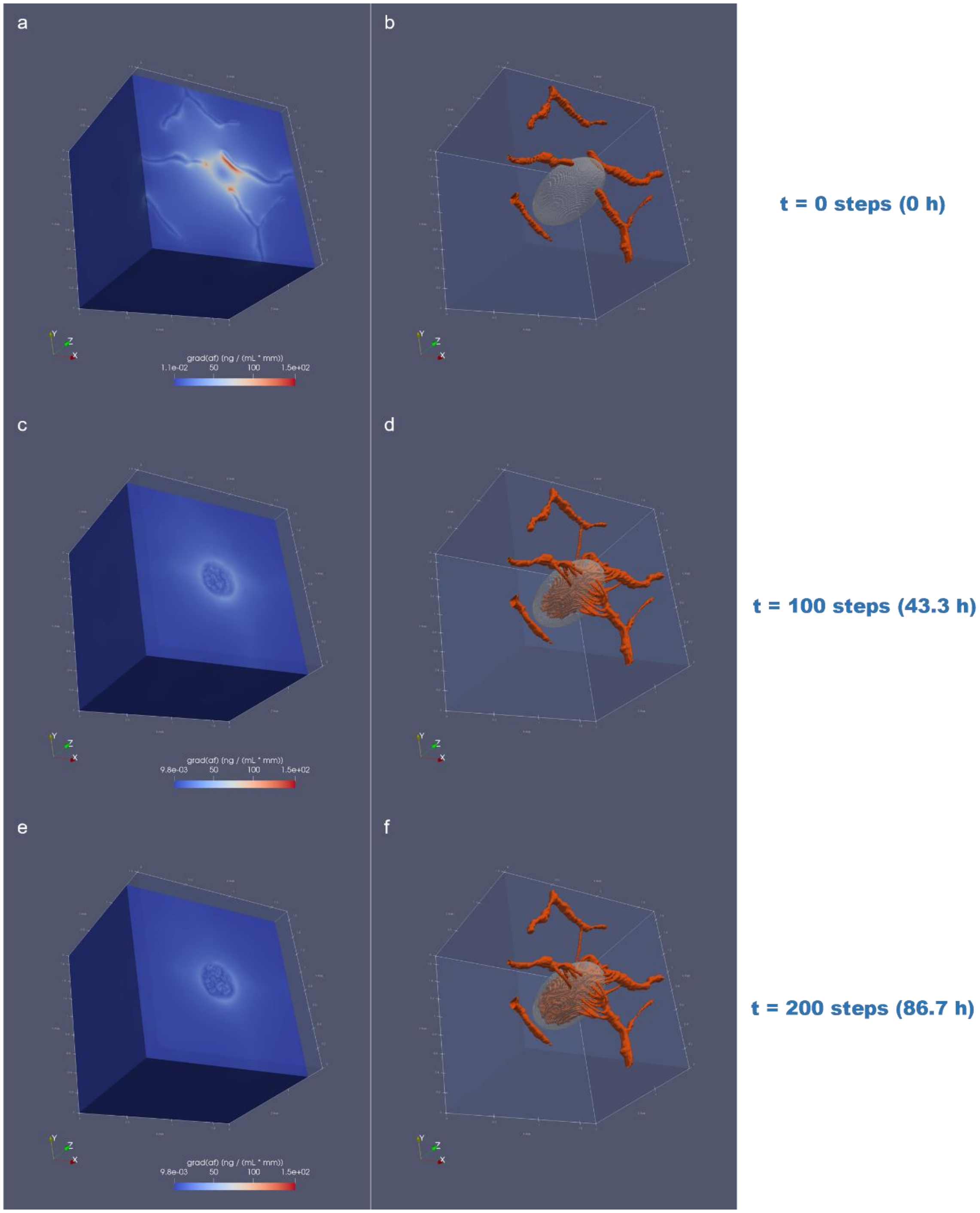


**Fig. S5.** AFs' gradient magnitude (a, c, e) while angiogenesis progresses (b, d, f) for the simulation reported in Figure 2 of the main text. In figure a, c, and e, different sections are shown. a) We evidence that the gradient is stronger near the capillaries, activating sprouting angiogenesis. c) and e) We evidence that the gradient is much lower, but still sufficient to drive tip cells toward the complete vascularization of the tumor. In general, the observed range for the magnitude of the AFs' gradient is [9.8e-3, 150] [ng / (mL * mm)]

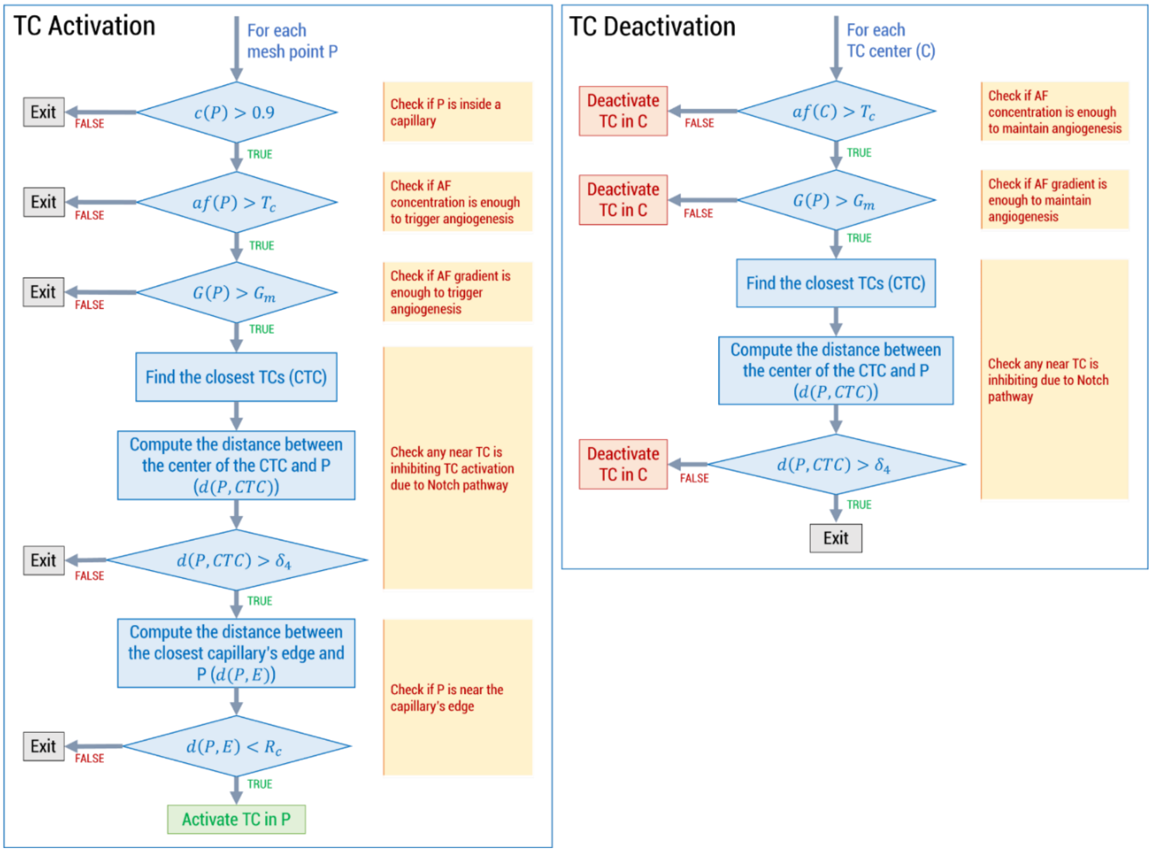


**Fig. S6.** Algorithms regulating TCs' activation (left) and deactivation (right).

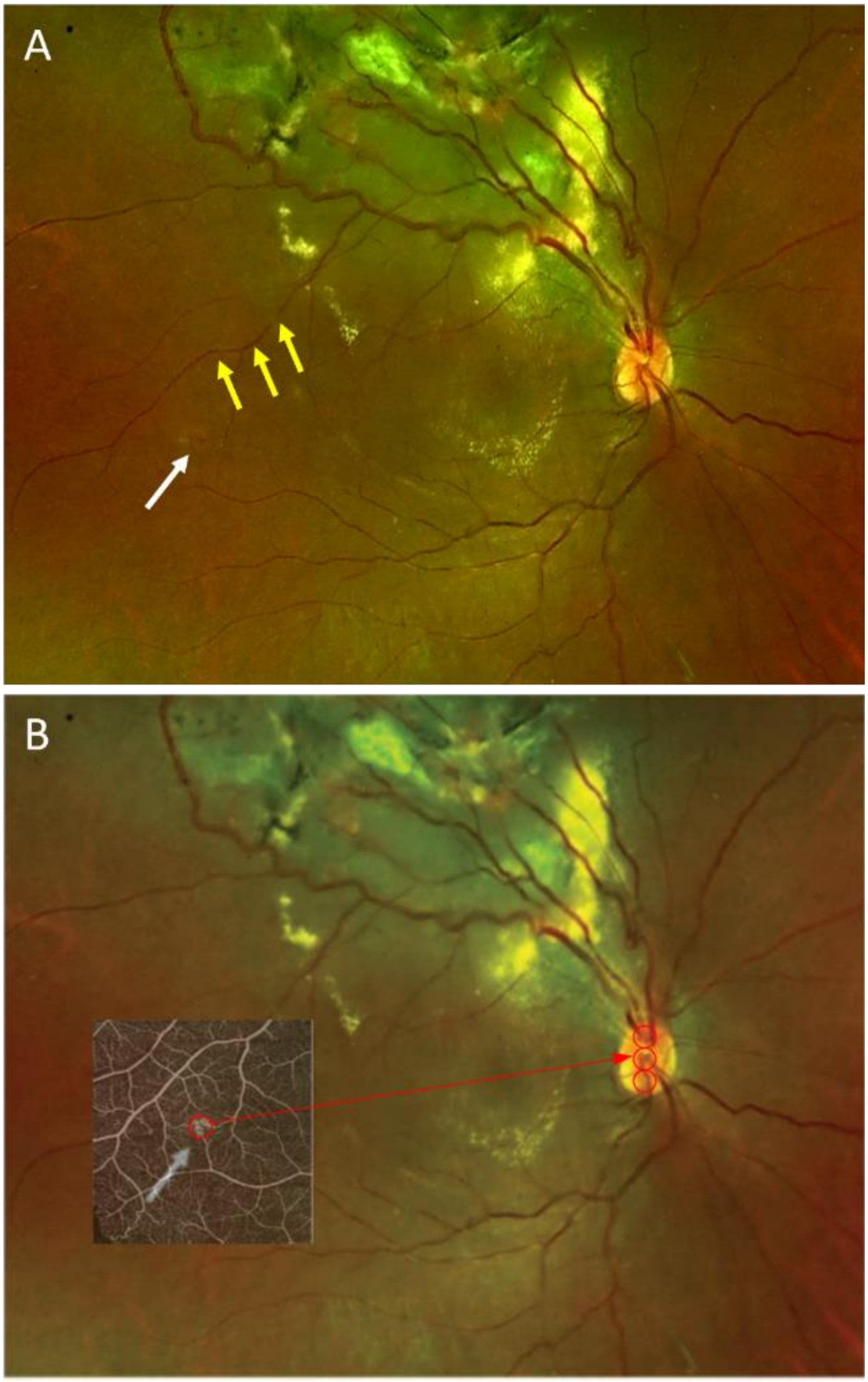


**Fig. S7.** Original fundus image (FI) of the lesion. The white arrow points to the RH, while the yellow arrows point to the arteriole that we used to align the FI with the OCTA image. B) Alignment between OCTA and FI, showing that the RH diameter is about 1/3 of the vertical OD axis.

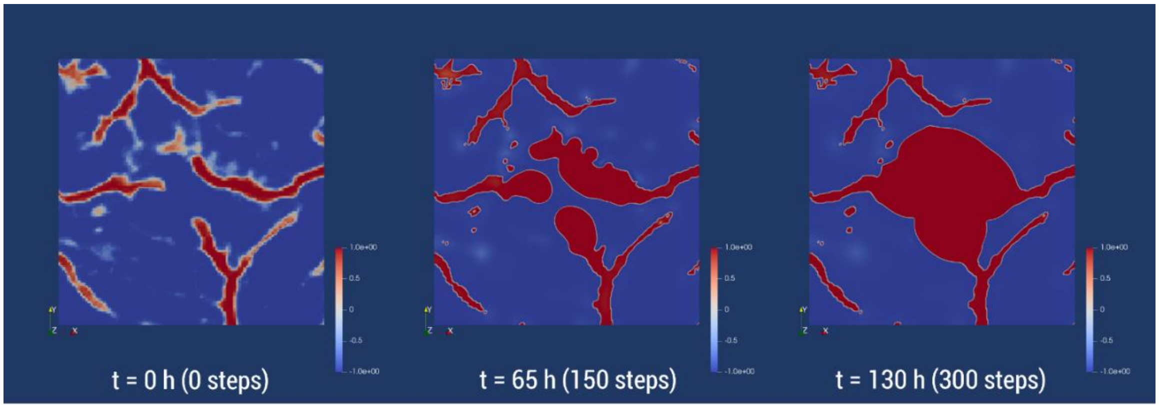


**Fig. S8**. Simulation in 2D with $\boldsymbol{\alpha_p = \alpha_{pSC}}$, showing major capillaries enlargement after 130 hrs (300 steps). Since in the selected case report we cannot see any sign of enlargement, the model does not reproduce reality for this parameter choice.

**Fig. S9.** Construction of the putative initial capillaries network (PICN). Image A is the original

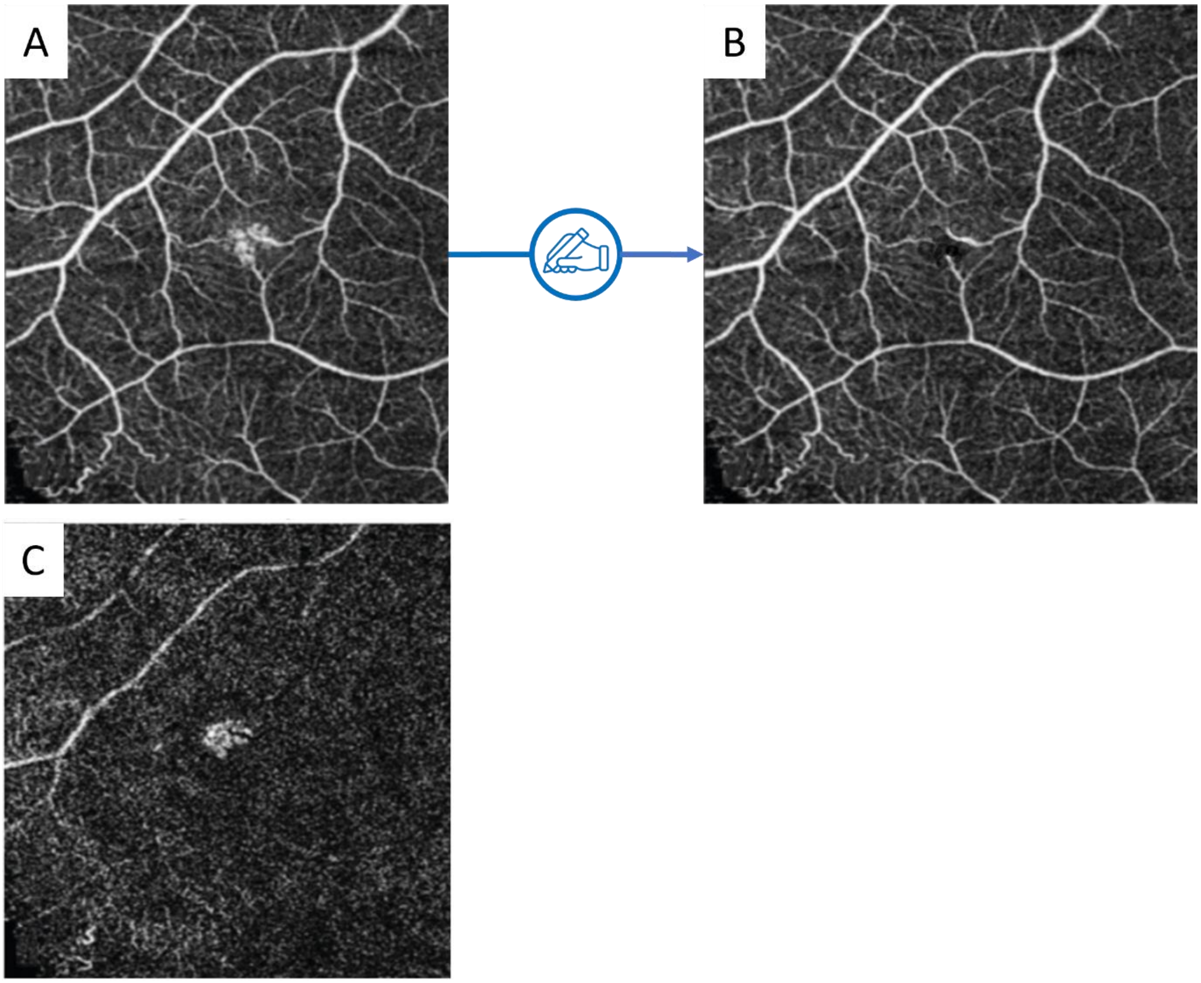


obtained by Goswami et al. (2021) with superficial OCTA. Image B was obtained by manual modification of A, using C as a reference to find the capillaries induced by the tumor.

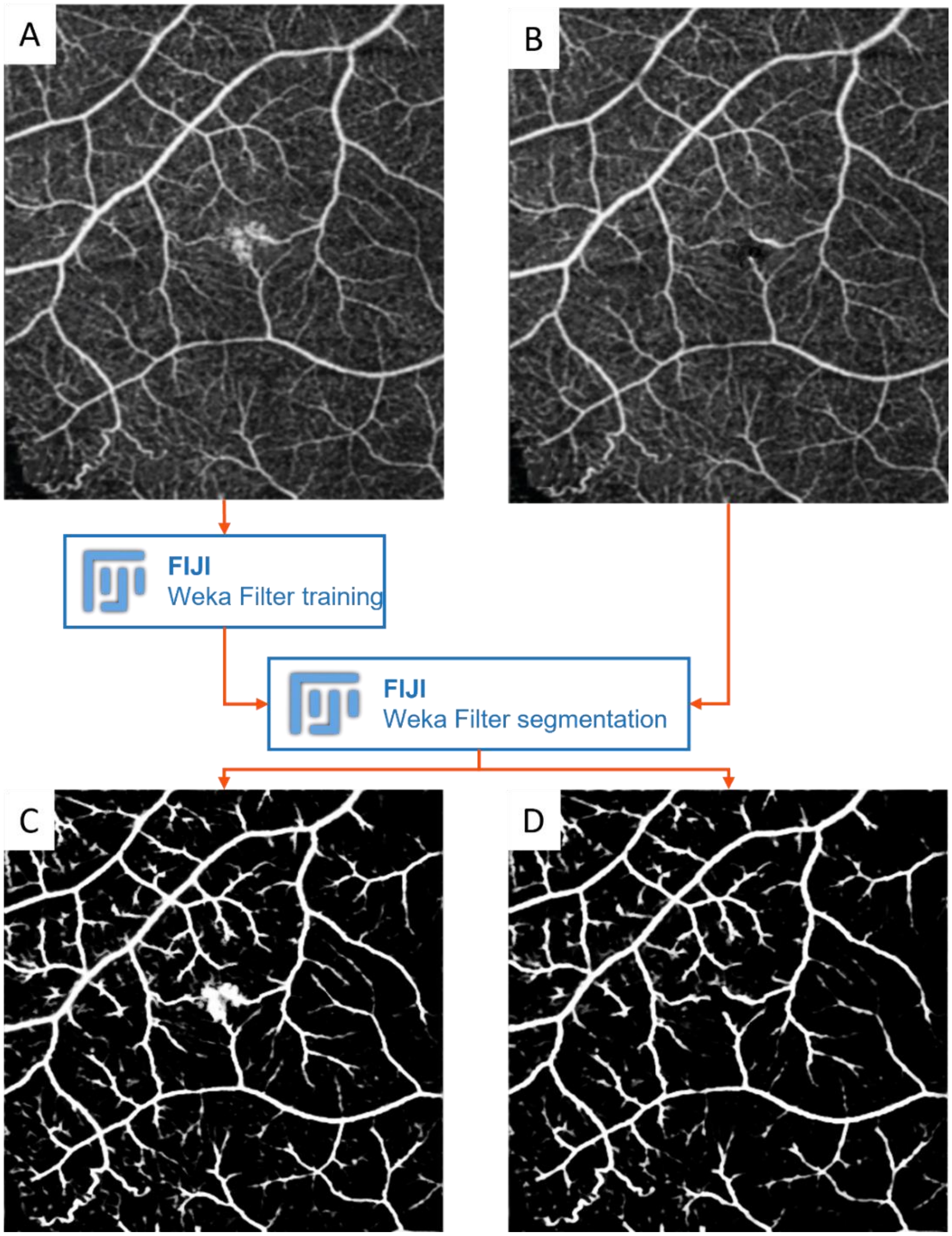

**Fig. S10.** Result of Weka segmentation on the original images.

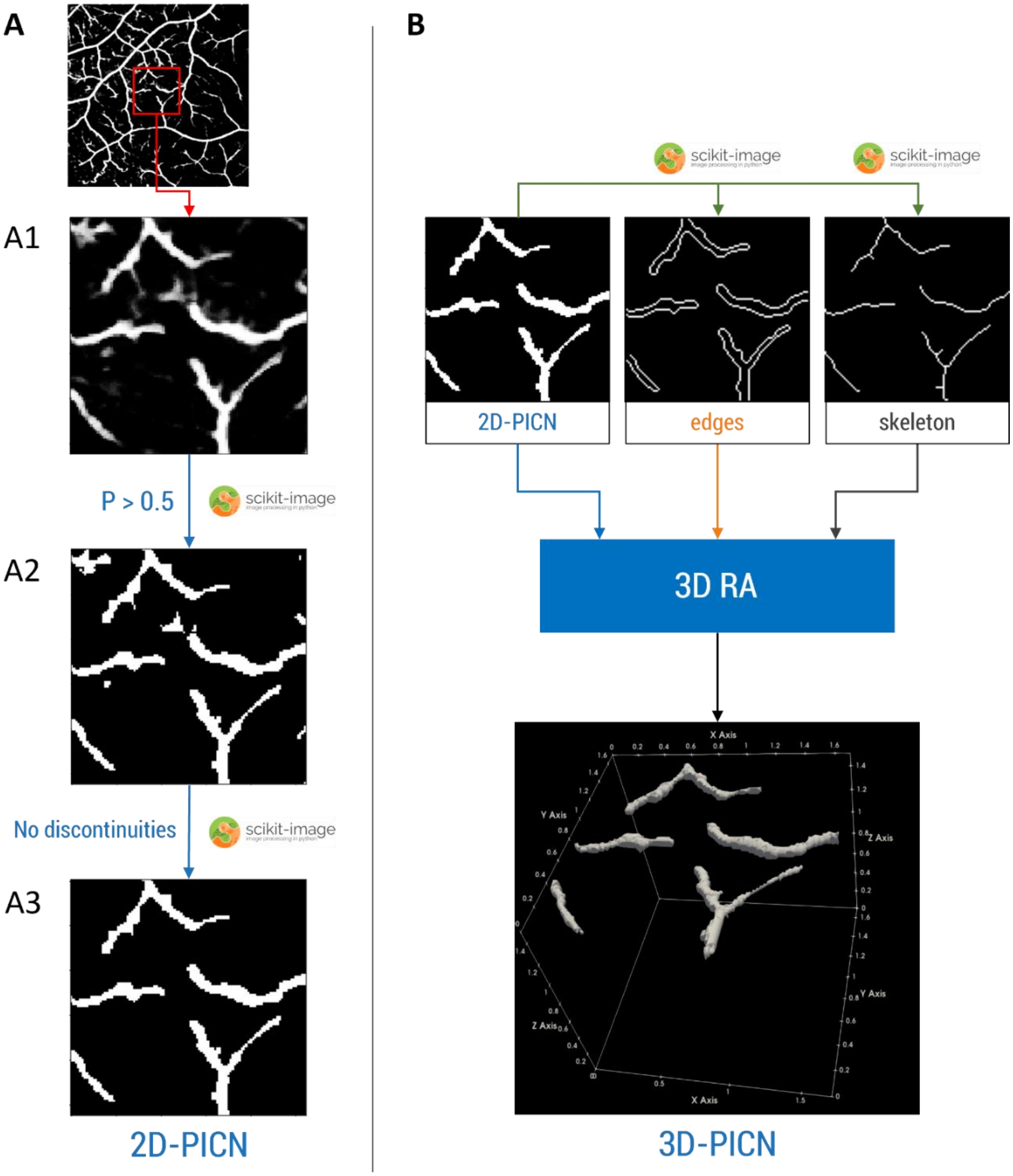


**Fig. S11.** A) image processing steps using Scikit-image. B) inputs and output of the 3D reconstruction algorithm (RA).

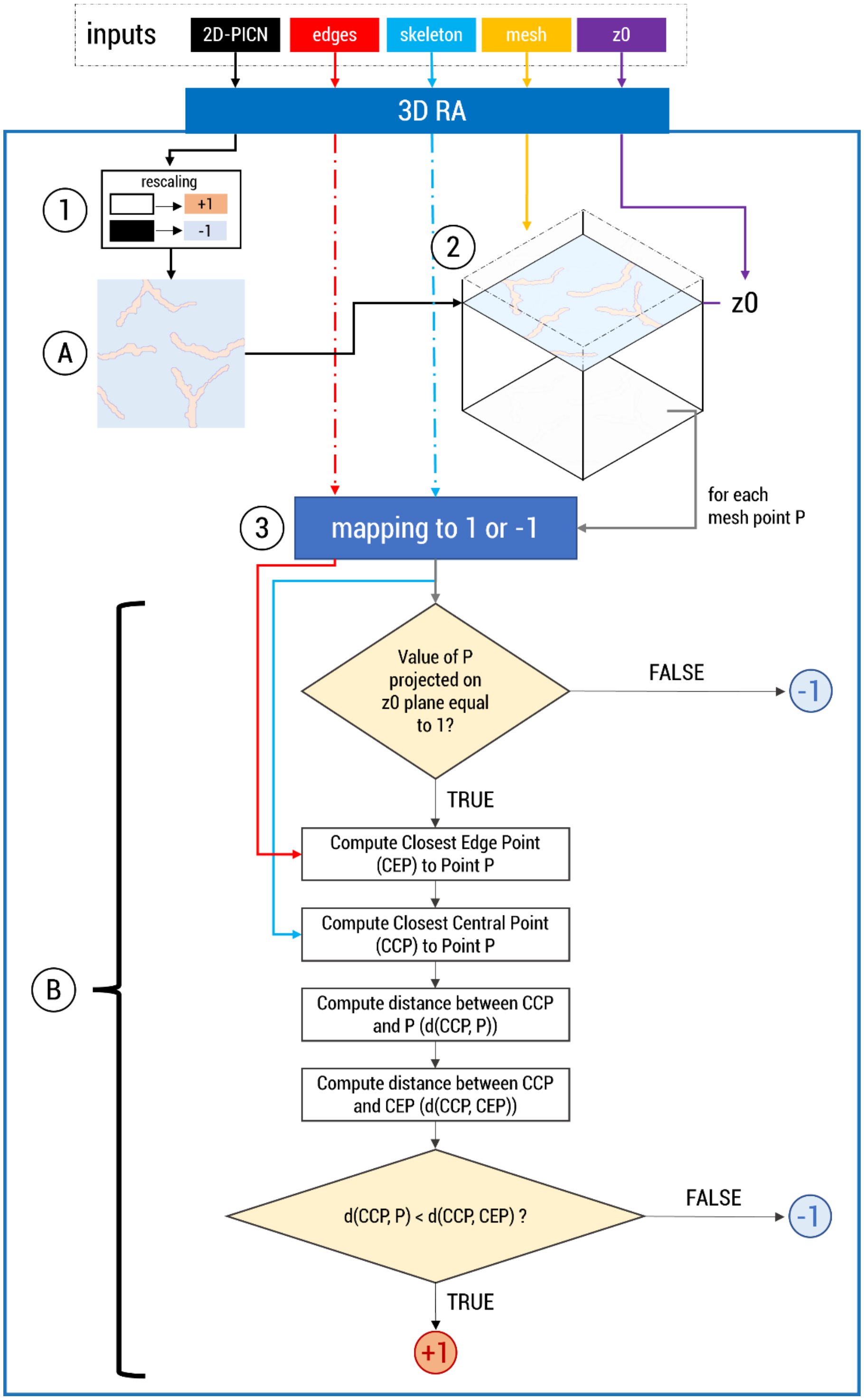


**Fig. S12.** Schematic representation of our 3D RA. On the top, we represented the inputs of the algorithms. (1) The first step rescales the 2D-PICN to obtain a 2D scalar field equal to 1 inside the capillaries and -1 otherwise (A). (2) The values of Matrix A are then mapped on the z0 plane of the 3D mesh. The user must specify the value of z0. (3) all other points are then mapped to 1 or -1 according to the procedure shown in the example. Notice that the algorithm employs the edges and the skeleton to compute the edge and center points.

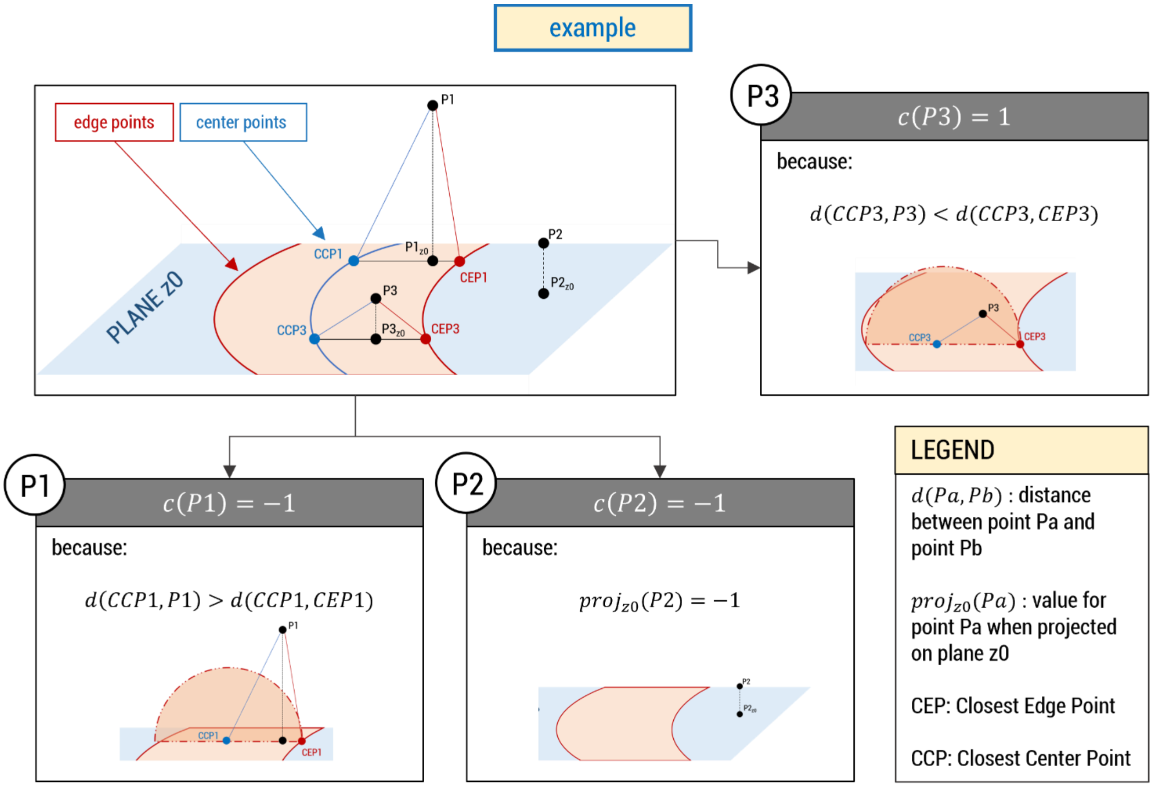


**Fig S13.** Examples of the application of the 3D mapping (Fig. S12B) for 3 points (P1, P2, and P3).

| $D_{af}$ value | Reference |
|---|---|
| $\mathbf{10^{-4}}\left[\frac{\boldsymbol{mm^2}}{\boldsymbol{s}}\right]$ | (Lai & Friedman, 2019) |
| $\mathbf{1.16 \cdot 10^{-6}}\left[\frac{\boldsymbol{mm^2}}{\boldsymbol{s}}\right]$ | (Guerra et al., 2021) |
| $\mathbf{2.22 \cdot 10^{-6}}\left[\frac{\boldsymbol{mm^2}}{\boldsymbol{s}}\right]$ | (Phillips et al., 2020) |
| $\mathbf{10^{-7}}\left[\frac{\boldsymbol{mm^2}}{\boldsymbol{s}}\right]$ | (Levine et al., 2001) |

**Table S1**. Some estimations of VEGF diffusivity reported in the literature.

**Movie S1 (separate file)**. Video of the simulation reported in Figure 2 of the main text. The capillaries are plotted every 10 time steps. On the left we show a 3d rendering of the occurring vascularization, while on the right we show the two sections reported in Figure 2.
